# Phase-shifting mid-infrared optothermal microscopy for wide-field hyperspectral imaging of living cells

**DOI:** 10.1101/2023.06.17.545432

**Authors:** Tao Yuan, Lucas Riobo, Francesca Gasparin, Vasilis Ntziachristos, Miguel A. Pleitez

## Abstract

Fast live-cell hyperspectral imaging at large field-of-views (**FOVs**) and high cell confluency remains challenging in vibrational microscopy due to the need for point-by-point focal excitation scanning. Imaging at high cell confluency and large FOVs is important, respectively, for proper cell function and statistical significance of measurements. Here, we introduce phase-shifting mid-infrared optothermal microscopy (**PSOM**) which interprets molecular-vibrational information as the optical path difference (**OPD**) induced by mid-infrared absorption and is capable of taking snapshot vibrational images over broad mid-infrared excitation areas at high live-cell confluency. By means of phase-shifting, PSOM suppresses noise to a quarter of current optothermal microscopy modalities to allow capturing live-cell vibrational images at FOVs up to 50 times larger than state-of-the-art. Additionally, it reduces illumination power flux density (**PFD**) down to 5 orders of magnitude lower than conventional vibrational microscopy—thus, considerably decreasing the possibility of cell photodamage.

## Introduction

Vibrational spectroscopy-based microscopy methods, such as coherent anti-Stokes Raman scattering (**CARS**),^1, 2^ stimulated Raman scattering (**SRS**),^3^ and mid-infrared (**mid-IR**) microscopy, are label-free imaging modalities that take advantage of the bond-specific vibrational transitions of biomolecules to achieve intrinsic chemical-contrast in living cells and tissues. In particular, in vibrational microscopy, hyperspectral images (where each pixel comprises a broad vibrational spectrum) can be acquired to allow the differentiation of biomolecule sub-groups—i.e. to discriminate among different types of lipids, proteins and carbohydrates by computational analysis of hyperspectral images.^4, 5^ In this way, label-free chemical microscopy could be applied for live-cell metabolic imaging—avoiding the use of external fluorescent labels needed in conventional florescence microscopy, which can undergo photobleaching,^6^ photoconversion,^7^ and possibly alter the metabolic activity of cells.^8^ However, most vibrational imaging methods apply tightly focused excitation and point-by-point raster-scanning for image formation, which results in slow spectral-imaging speed (minutes to hours per wavelength in mechanical scanning) precluding acquisition of live-cell hyperspectral images and increasing the risk of photodamage of living cells.^9, 10^

To increase imaging speed—profiting from mid-IR absorption with cross sections eight orders of magnitude larger than Raman scattering—wide-field mid-IR microscopy methods based on optothermal effect and optical phase-contrast detection for fast hyperspectral imaging have been recently proposed. For instance, Cheng *et al*. developed bond-selective transient phase (**BSTP**) imaging, in which mid-IR excitation results in a transient refractive index change in the excited sample that is detected by a phase-contrast microscope. BSTP enabled hyperspectral imaging in living cells at above-video-rate speed, i.e., above 24 frames per second (**fps**).^12^ Similarly, Ideguchi *et al*. developed molecular vibration-sensitive quantitative phase-contrast imaging (**MV-QPI**), which detects mid-IR absorption images by quantitative phase-contrast detection.^13^ Just like BSTP, MV-QPI has also shown above-video-rate label-free mid-IR hyperspectral imaging in living HeLa cells with sub-cellular spatial resolution. However, despite offering high-imaging speed and demonstrations of their feasibility for live-cell hyperspectral imaging BSTP and MV-QPI are restricted to small fields-of-view (FOVs) of 10–50 μm, whereas FOVs above 100 μm are necessary for imaging cell populations. To address the limitation in FOV size, we recently introduced wide-field optothermal mid-infrared microscopy (**WOMiM**) based on snapshot pump-probe detection of optothermal signal.^14^ With WOMiM, we demonstrated chemical-contrast imaging of an unprecedentedly wide FOV of up to 180 μm, achieving hyperspectral imaging of lipids (triglycerides) in the 2950 to 2830 cm^-1^ range. However, due to low sensitivity, the working principle of WOMiM was demonstrated only on synthetic phantoms of triglyceride drops.

In conventional wide-field optothermal imaging, phase difference (termed MIR-phase) micrographs induced by mid-IR absorption are obtained by subtracting a strong phase background (termed intrinsic-phase) caused by a sample’s intrinsic optical properties (e.g., refractive index). The quantitative intrinsic-phase image is usually removed using postprocessing methods such as, Hilbert transform,^11^ derivative method,^12^ and non-linear phase unwrapping method.^13^ However, since the MIR-phase is two to three orders of magnitude smaller than the intrinsic-phase,^14, 15^ these postprocessing methods usually alter the weak MIR-phase signal, resulting in poor detection sensitivity. Although adaptive dynamic range quantitative phase imaging (QPI) has been recently developed to bridge the large phase gap between intrinsic-phase and MIR-phase,^15^ the strong probe illumination light needed for high sensitivity phase detection increases the risk of phototoxicity to the cells, and large FOVs for imaging multiple living cells have not been demonstrated. As a step forward in wide-field mid-IR optothermal microscopy, we introduce here phase-shifting optothermal microscopy (**PSOM**), which implements wide-field excitation for a large FOV and incorporates a polarization-based phase-shifting module **(PSM)** to obtain chemical-contrast images with high sensitivity despite large intrinsic-phases. We hypothesized that achieving intrinsic-phase independent optothermal imaging will provide high sensitivity to small MIR-phase variations, thus allowing further expansion of the mid-IR beam for a larger FOV, and thereby reducing the mid-IR phototoxicity to cells. In this manuscript, we demonstrate how the phase-shifting module enables PSOM to reach shot-noise limited and effectively reduce noise-equivalent phase down to 0.26 mrad—which is a quarter of the state-of-the-art, obtained only under the extreme condition of strong probe irradiation.^15^ Low noise allows PSOM to sense a small MIR-phase variation under a wide-field and low power flux density (**PFD**) excitation. In our operation examples, PSOM achieved an unprecedentedly large FOV (50 times larger in area than state-of-the-art FOV)^16^ for live-cell mid-IR optothermal imaging, using mid-IR PFD 5 orders of magnitude below excitation PFD typically used in other vibrational imaging modalities, such as CARS.^17^ As examples of operation, we performed measurements of lipid droplets (**LDs**) in matured adipocytes at high confluence (100%), which are unprecedented in wide-field photothermal-microscopy due to strong intrinsic-phase of highly confluent cells.

## Results

### Basic working principle of PSOM

**Fig. 1** schematically depicts the working principle of pump-and-probe PSOM. PSOM is a label-free bond-selective imaging method based on wide-field mid-infrared excitation for optothermal generation and polarization-based phase-shifting for thermo-optic read-out (see **Fig. 1a**, more details in **Methods**). Unlike previous mid-IR wide-field optothermal (photothermal) microscopy methods,^14, 15, 18^ PSOM suppresses the large intrinsic-phase of the sample by means of a unique combination of polarization phase-shifting mechanism^19^ (see **Fig. 1b-f**) and MIR-phase retrieve algorithm, thus enhance chemical contrast sensitivity for live-cell imaging at high cell confluence and for capturing large FOVs. A detailed description of the system is presented in **Methods**. Here, briefly, a 532 nm linearly polarized (at 45° in relation to the x axis) probe-beam is split by a birefringent beam displacer into two orthogonally polarized beams, termed sample beam (**SB**) and reference beam (**RB**), which are subsequently rotated 90° by a half-wave plate (see **Fig. 1a**). The two parallel beams then travel through the sample plane and are recombined by a birefringent beam combiner (**Fig. 1b**), forming a balanced path Mach–Zehnder interferometer to cancel out the phase noise and reach shot noise limit.^20^ Depending on the optical phase difference between the SB and RB, the resulting recombined beam is either linearly (same phase) or elliptically (different phase) polarized (see **Fig. 1c-e**). For the sake of imaging, the recombined beam is collected by a microscope objective with its focus on the sample plane. The probe beam then travels through a quarter-wave plate (fast axis at 45°) resulting in a linearly polarized beam with a polarization angle that linearly depends on the optical phase difference between SB and RB (see **Fig. 1f**). The probe beam keeps its initial polarization angle (*α*, at 45°) when there is no phase difference between SB and RB, while it rotates to *β* when an optical phase difference δ_*0*_ between SB and RB is introduced (**Fig. 1f**)—for instance, on SB at the sample plane (**Fig. 1b**). After passing through an analyzer (i.e., a linear polarizer) and tube lens, the probe beam then forms an image (PSM image) on the camera sensor, where the intensity of PSM image correlates sinusoidally to the optical phase difference δ_0_ between SB and RB (see **Supp. Fig. 1** and **Supp. Fig. 2**). PSM images at different analyzer angles (*θ*) are acquired in order to determine the introduced optical phase difference δ_0_ (see **Supplementary § 1**).

**Fig. 1.**
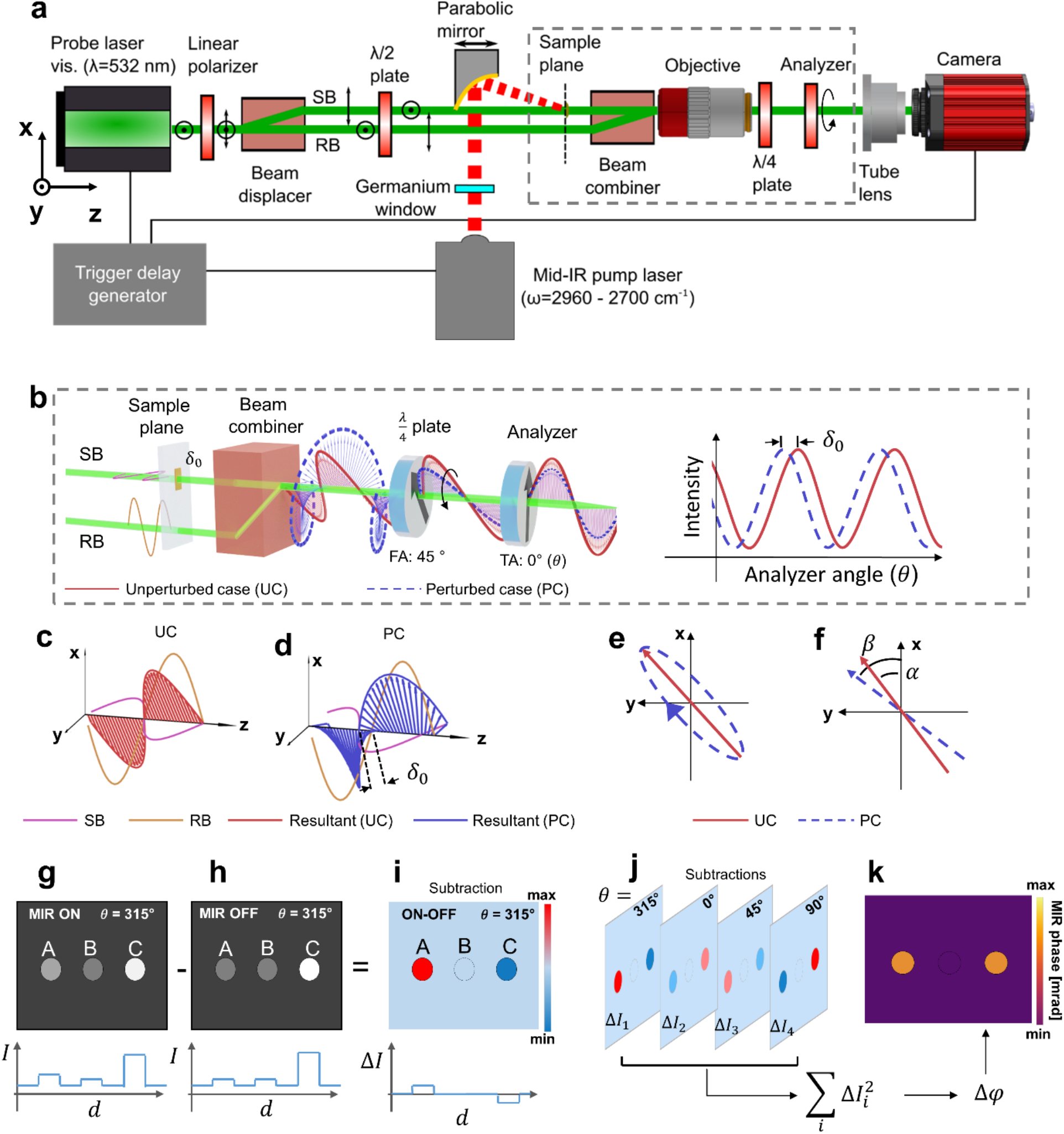
Imaging principle of phase-shifting optothermal microscopy (PSOM). **a**, Diagram of the overall system. PSOM is a synchronized pump-and-probe system where the phase-shifting module (PSM) probes the phase difference resulting from a wide-field mid-IR excitation (pump), to obtain a chemical contrast image. SB: sample beam; RB: reference beam. **b**, A phase perturbation *δ*_0_ in the sample plane results in elliptically polarized light after the beam combiner, and afterward, causes rotation of the polarization plane of the beam after the quarter-wave plate. The rotation of polarization plane is analyzed by changing the transmission axis (TA) of analyzer. FA: fast axis. **c**,**d**, Polarization states of the beams after the beam combiner. (c) Linear polarization is constructed in unperturbed case (UC), and (d) elliptical polarization is constructed in perturbed case (PC). **e**, Polarization states of (c) and (d) illustrated in x-y plane. **f**, Polarization states of the beam after the *λ*/4 plate. *α*: Polarization angle in unperturbed case; *β*: Polarization angle in perturbed case. **g-i**, Demonstration of MIR-ON and MIR-OFF subtraction at an analyzer angle of 315°, with line profile crossing three targets. A: absorption target; B: non-absorption target; C: absorption target at high dynamic range. I: intensity; d: distance. **j**, 4 subtraction images are obtained at 4 analyzer angles (*θ* = 315°, 0°, 45°, 90°). **k**, A PSOM image constructed by the square sum 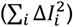 of the subtraction frames illustrates quantitative phase perturbation (∆*φ*) induced by mid-IR excitation, regardless of dynamic range of intrinsic-phase.

In PSOM, as see in **Fig. 1a**, a sample placed in the SB region of the sample plane is illuminated (wide-field illumination, see **Supp. Fig. 3**) by a pulsed mid-IR laser beam, at a selected excitation wavenumber, in order to generate optical phase perturbation according to molecular vibrational absorption. PSM images are then sequentially acquired at optothermal transition (MIR-ON) and at thermal relaxation between mid-IR laser pulses (MIR-OFF, see **Methods** and **Supp. Fig. 4**) to generate a subtraction image (MIR-ON minus MIR-OFF, see **Fig. 1g-i**). Four subtraction images at four analyzer angles (namely: 315°, 0°, 45°, and 90°) are needed for phase-shifting mechanism (**Fig. 1j**). A PSOM micrograph is then obtained by taking the square sum of the subtraction images and applying equation S11 (abbreviated as: eq. S11, see **Supplementary § 2**) to generate a phase map (**Fig. 1k**), which linearly correlates with the sample’s mid-IR absorption at the excitation wavenumber (see **Supplementary § 3**).^14^ As detailed in **Supplementary § 2**, the square sum of the subtraction images suppresses the background from sample’s intrinsic-phase, resulting in enhanced MIR-phase contrast images that exclusively relate to mid-IR absorption.

### System characterization and proof of concept

First, we tested and characterized the imaging and spectroscopic abilities of PSOM using synthetic triglyceride (**TAG**) phantoms (see **Methods**). **Fig. 2a**, for example, depicts a PSOM micrograph of TAG drops in water acquired at a mid-IR excitation wavenumber of 2850 cm^-1^ (3.509 μm), assigned to symmetric CH^2^ vibration and generates intrinsic molecular contrast for lipids at a maximum MIR-phase of 81.5 mrad. For comparison, **Fig. 2b** shows a PSOM micrograph of the same FOV with no mid-IR excitation, which results in an image where—as expected—no structures can be observed and from which a spatial noise-equivalent phase of 0.33 mrad can be determined. Furthermore, continuous MIR-OFF measurement of 57 minutes obtained temporal noise-equivalent phase of 0.26 mrad (see **Supp. Fig. 5**, for calculation of noise-equivalent phase, See **Methods**). Such a low temporal noise level is a quarter of the values reported for state-of-the-art optothermal microscopy^15^ and yielded a contrast-to-noise ratio (**CNR**, see **Methods**) of 282:1 for the marked TAG drop in **Fig. 2a**. Interestingly, when comparing the imaging resolution of PSOM (**Fig. 2a**) and PSM (intensity image, **Fig. 2c**) for the same structure, PSOM image appeared sharper than PSM image despite having been generated by mid-IR excitation, which suggests that PSOM might result in higher imaging resolution compared to PSM. This is demonstrated in **Fig. 2d** where the intensity profiles across the smallest TAG drop observed in **Fig. 2a**,**c** (white dashed line) were compared—the full-width-at-half-maximum (**FWHM**) of PSOM profile was 2.0 μm while the FWHM of PSM’s profile was 3.1 μm. The smaller FWHM value from PSOM could have resulted from the fast thermal dissipation around the interface of TAG drops and water; as noted by other authors using other wide-field optothermal modalities; For instance, BSTP imaging.^14^ For reference, **Fig. 2d** also shows the MIR-phase profile for the MIR-OFF micrograph in **Fig. 2b**, where the signal and noise level obtained with PSOM are compared.

**Fig. 2.**
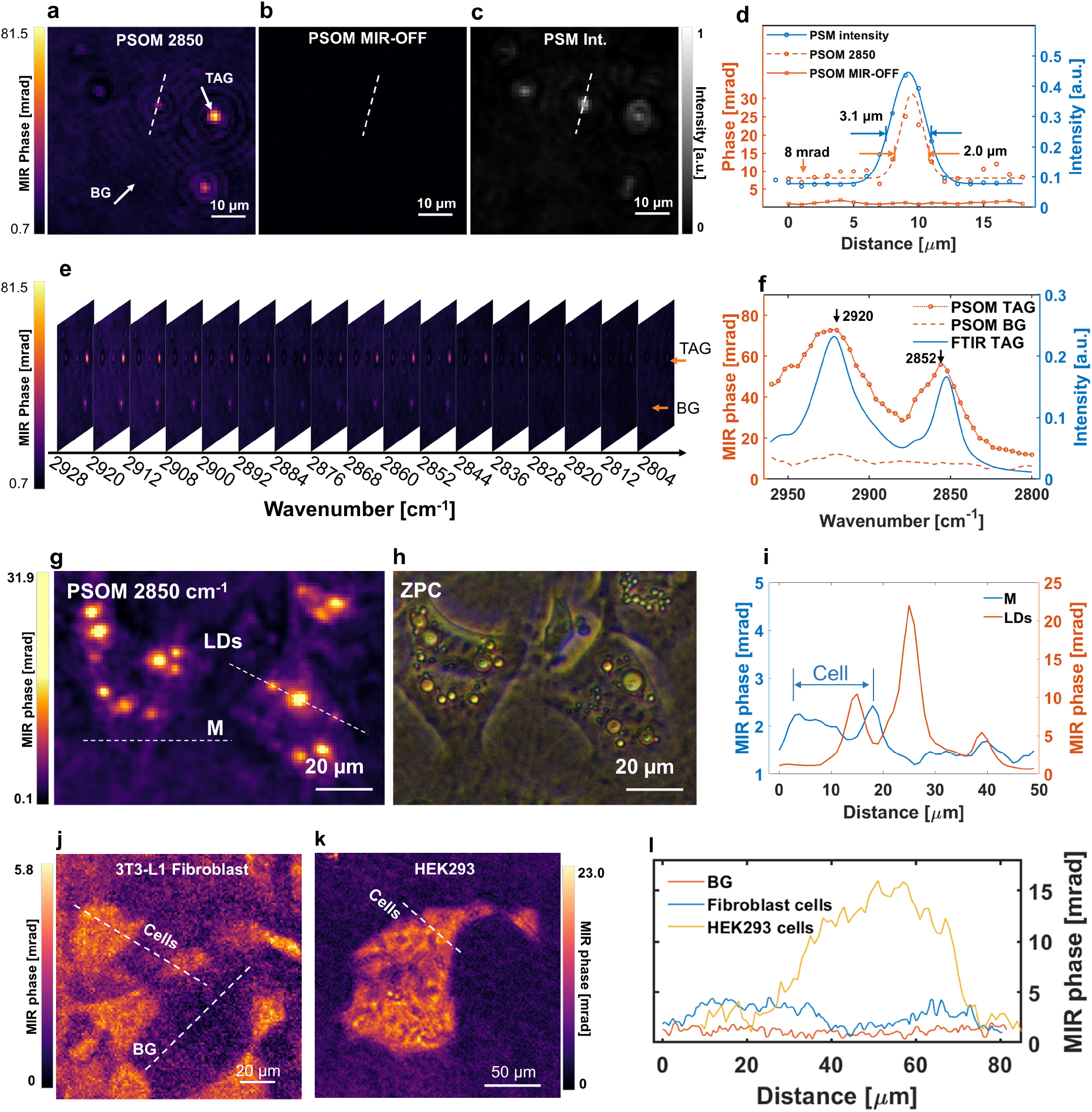
Imaging system validation in synthetic triglyceride (TAG) drops and living cells. **a**,**b**, PSOM micrographs of the triglyceride (TAG) drops (a) with mid-IR excitation at wavenumber 2850 cm^-1^, and (b) without mid-IR excitation. BG: background. **c**, Intensity image acquired by phase-shifting module (PSM), at an analyzer angle of 45°. **d**, Line profiles crossing the smallest TAG drop in (a-c). **e**, PSOM hyperspectral image stack of the same FOV as (a). **f**, PSOM spectrum of TAG drop extracted from (e) and its comparison to FTIR measurement. **g**, PSOM micrograph of single living cells (3T3-L1), demonstrating CH_2_ vibrational contrast obtained at excitation wavenumber of 2850 cm^-1^. Dotted lines drawn through lipid droplets (LD) and plasma membrane (M). **h**, The same FOV as (g) acquired by Zernike Phase contrast (ZPC) microscope. **i**, Line profiles crossing lipid droplets (LDs) and the membrane (M) of a cell, as indicated in (g). **j**,**k**, PSOM image of 3T3-L1 fibroblast cells (j), and a group of HEK293 cells (k) acquired at mid-IR wavenumber of 2850 cm^-1^. **l**, Line profiles of the cells from (j) and (k).

Beyond single-wavelength imaging, hyperspectral imaging (**Fig. 2e**) was achieved for the same FOV in **Fig. 2a** by acquiring PSOM images while tuning the excitation wavenumber along a mid-IR spectral range from 2960 to 2800 cm^-1^ in steps of 4 cm^-1^ (for simplicity, only 17 out of 41 frames are shown). This allowed us to selectively extract the spectrum of structures of interest in the FOV; as demonstrated, when we plotted the spectrum (**Fig. 2f**) for the structure marked in **Fig. 2a** (white arrows). Clearly, from **Fig. 2f**, the spectrum acquired by PSOM showed two absorption peaks characteristic of TAG at 2920 and 2852 cm^-1^, attributed to methylene’s (CH^2^) asymmetric and symmetric stretching vibration, and in good agreement with spectrum of TAG obtained by conventional Fourier-transform infrared spectroscopy (**FTIR**). The MIR-phase at 2920 and 2852 cm^-1^ was up to 72.7 mrad and 52.8 mrad, respectively; while, for reference, the spectrum extracted from background (**BG**), originating from water absorption, remained low at a mean MIR-phase intensity value of 8 mrad. The acquisition time of this hyperspectral imaging stack was 32.8 s (discarding data transferring time, see **Methods** for details). Importantly, experiments in synthetic TAG phantoms demonstrated linear response of MIR-phase intensity with mid-IR excitation power (**Supp. Fig. 6**), revealing PSOM’s ability to detect TAG drops at an excitation PFD as low as 0.05 µW/µm^2^ at CNR of 26:1. However, in order to enhance sensitivity in the experiments presented here, we used PFD excitation of 0.16 *µ*W/µm^2^, which is still 3 times lower than typical values reported for other photothermal imaging modalities^16^ (see **Supp. Table 1**) and up to 4 orders of magnitude below typical excitation values applied in other vibrational imaging modalities, such as CARS (15 mW/*µ*m^2^).^17^

The exceptionally low mid-IR PFD used in PSOM (0.16 *µ*W/*µ*m^2^) facilitates the application of mid-IR optothermal microscopy to the study of live-cell metabolism by minimizing phototoxicity exerted on cells (viability test results are shown in **Supp. Fig. 7**, viability of HeLa cells was 85.4% after 2 hours of measurement with PSOM). A demonstration of PSOM’s live-cell imaging ability is presented in **Fig. 2g** for 3T3-L1 differentiated adipocytes, showing lipid contrast obtained at 2850 cm^-1^ (CNR 117:1) generated by lipid droplets (LDs) formed after 6 days of incubation (see **Supp. Fig. 8**,**9** for corresponding PSM intensity and intrinsic-phase images). The PSOM micrograph in **Fig. 2g** is comparable to the image obtained by conventional Zernike phase contrast (**ZPC**) microscopy as shown in **Fig. 2h**, indicating good morphological correlation between the two techniques. The MIR-phase of lipid droplets ranges from around 5 mrad to 22 mrad while contrast from the phospholipids in the intracellular cell membranes at 2850 cm^-1^ peaks at 2.4 mrad (**Fig. 2i**). Such a low MIR-phase contrast range (2.4 – 22 mrad) from live-cell structures, at least three orders of magnitude smaller than the intrinsic sample phase^25,26^ (c.a. 19 rad, see **Supp. Fig. 10)**, is hard to be preserved in conventional QPI after application of post processing algorithms, typically applied to remove phase-wrapping.^11-13^ Thus, benefiting from intrinsic-phase removal and reduced noise, PSOM is able to detect low-contrast cellular structures, such as cytoplasmic and intracellular membranes—rich in CH_2_ bonds. **Fig. 2j**,**k** show PSOM images of 3T3-L1 fibroblast cells and HEK293 cells, targeting the CH_2_ vibrational signal using mid-IR excitation wavenumber of 2850 cm^-1^. **Fig. 2l** shows line intensity profiles crossing the two types of cells. It was observed that fibroblast cells with low confluence—sparse cell distribution—have MIR-phases of around 4 mrad for a single cell. In contrast, highly confluent HEK293 cells—which grow in dense groups—have MIR-phases that can reach up to around 15 mrad.

### Implementation of PSOM for large FOV live-cell mid-IR microscopy

**Fig. 3** depicts an operation example of large FOV imaging of living cells at high confluence (100%), which has not been demonstrated before by state-of-the-art optothermal microscopy. Measuring highly confluent live cells is important because many cellular processes are only representative of the *in vivo* state at high confluence; for instance, the differentiation of myoblasts into myotubes and the differentiation of preadipocytes into adipocytes.^21^ **Fig. 3a** shows a ZPC image of 3T3-L1 cells at day 6 after differentiation, displaying an imaging area fully occupied by cells (100% confluence)—for the same FOV, **Fig. 3b** illustrates a PSOM image at 2850 cm^-1^ with selective lipid contrast. The mid-IR irradiation area for PSOM imaging is an ellipse (major axis: 700 μm, minor axis: 400 μm) as shown by the white dashed line in **Fig. 3b**, giving an effective chemical-contrast FOV of 2.2×10^5^ μm^2^. Larger FOVs can be obtained by further expanding the mid-IR excitation area, as shown in **Supp. Fig. 11a** for matured 3T3-L1 differentiated cells in a FOV of 5.18×10^5^ *µ*m^2^—which is 50 times the demonstrated FOV of state-of-the-art wide-field optothermal microscopy (acquisition time: 0.8 s, discarding data transfer time). Such a large imaging area with living cells at 100% confluence is unprecedented in optothermal microscopy.

**Fig. 3.**
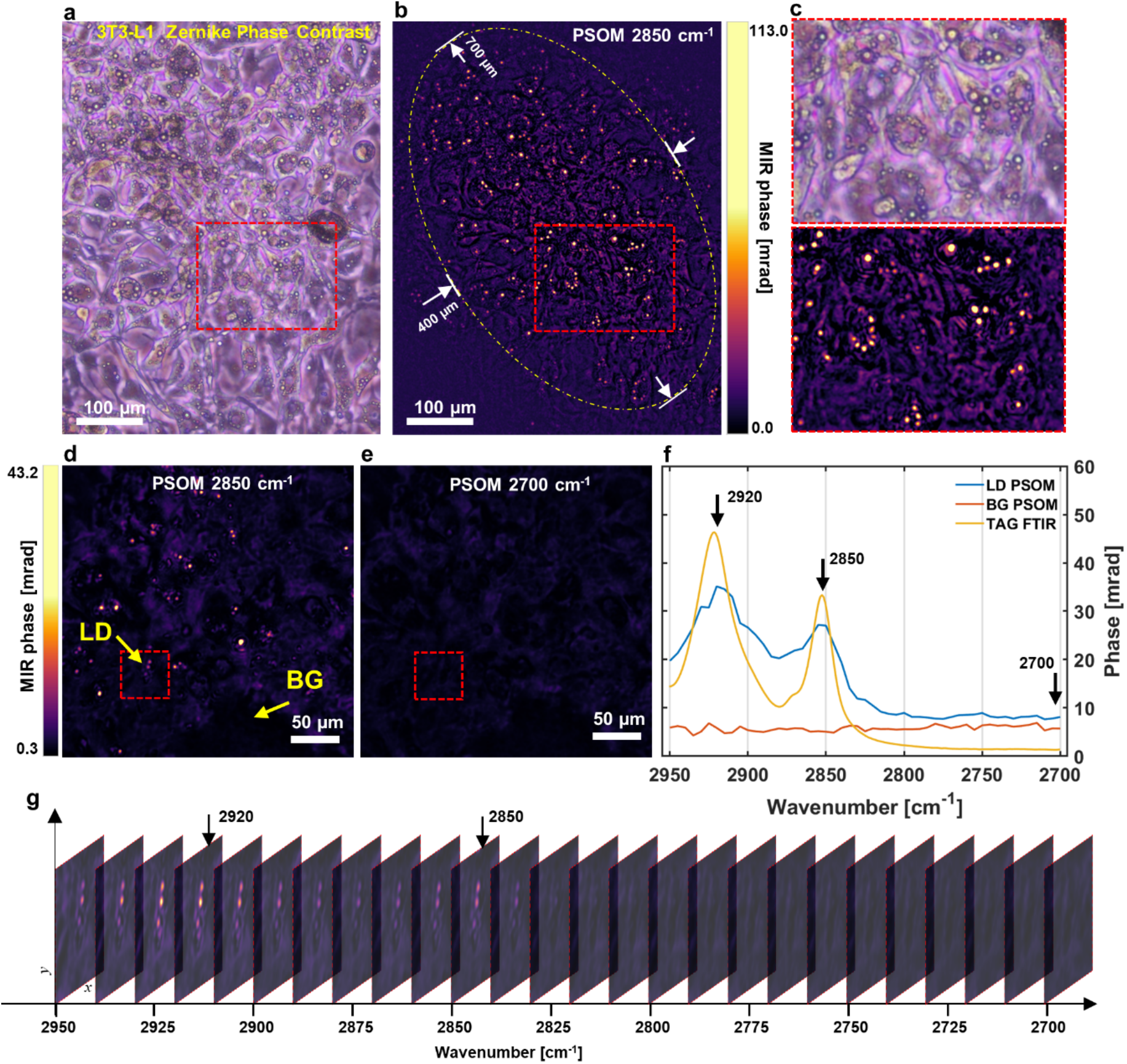
Highly confluent (100%) living cells visualized by chemical-contrast imaging and hyperspectral imaging using PSOM. **a**, 3T3-L1 cell micrograph (day 6 after differentiation) obtained by Zernike phase contrast microscope. **b**, Chemical-contrast micrograph obtained by PSOM for the same FOV as (a), at wavenumber 2850 cm^-1^. Dashed ellipse: effective chemical-contrast FOV of 400×700 um^2^. **c**, Zoomed in FOVs in red dashed rectangles in (a) and (b). **d**,**e**, PSOM images of 3T3-L1 cells at (d) CH_2_ resonance wavenumber 2850 cm^-1^, and (e) CH_2_ off-resonance wavenumber of 2700 cm^-1^. These are two representative images selected from a hyperspectral imaging stack. **f**, Spectra extracted from hyperspectral imaging stack, showing spectrum of lipid droplet and spectrum from background (cell media). The Fourier-transform infrared spectroscopy (FTIR) measurement of triglyceride (TAG) is plotted together for comparison. **g**, Hyperspectral imaging stack for a selected area marked by rectangles in (d) and (e), illustrating chemical-contrast change of three lipid droplets from wavenumber of 2950 cm^-1^ to 2700 cm^-1^. LD: lipid droplet; BG: background.

**Fig. 3c** shows a zoomed-in FOV of the red dashed area highlighted in **Fig. 3a** and **Fig. 3b**, used to further compare images from ZPC microscope and PSOM. In ZPC micrographs, such as in **Fig. 3a**,**c**, lipid droplets can only be recognized by overall morphology (i.e., with no specific lipid contrast), which makes them hard to distinguish from the large background originating from all molecular content in the cell body—in particular, by image segmentation algorithms (see **supp. Fig. 12**). Therefore, quantification of the amount of lipid droplets cannot be easily automated in conventional ZPC micrographs. This confounding background is effectively suppressed in PSOM micrographs (**Fig. 3b**,**c**), where the contrast at 2850 cm^-1^ originates predominantly from the lipid droplets. Unlike measurements at low confluence (such as the single-cell images in **Fig. 2g)**, the MIR-phase of lipid droplets in **Fig. 3b**,**c** reaches up to 113.0 mrad (CNR 404:1), possibly due to more efficient lipogenesis at 100% confluence as opposed to lower confluence values. Moreover, acquiring the intrinsic-phase of highly confluent cell populations by conventional QPI methods is prone to errors due to strong phase wrapping (see **Supp. Fig. 10**), which makes conventional QPI-based wide-field optothermal microscopies inadequate for the task of imaging live-cell populations at high confluence. PSOM circumvents this problem by acquiring native MIR-phase images—i.e., independent of intrinsic-phase. The capability to image large FOVs enables PSOM to simultaneously capture the heterogeneous activity of cells at different locations, and hence allows statistical analyses to be carried out on large number of cells to avoid biases that arise from analyzing single cells. As an example, we applied PSOM to determine LD count and LD displacement in a large FOV and at high confluency taking advantage of lipid-specific contrast, see **Supp. Fig. 11b-e** and **Supp. Fig. 12**.

Finally, **Fig. 3d**,**e** show two PSOM micrographs of 3T3-L1 cells, at lipid-specific mid-IR wavenumber of 2850 cm^-1^ and lipid unspecific (off-resonance) wavenumber of 2700 cm^-1^, used to demonstrate PSOM image specificity to molecular absorption when visualizing living cells. Comparing **Fig. 3d**, and **Fig. 3e**, it can be observed that the lipid droplets only appear when a wavenumber of 2850 cm^-1^ was used, and are absent when a wavenumber of 2700 cm^-1^ was utilized. Moreover, **Fig. 3f** shows the spectrum of the marked LD in **Fig. 3d**, as well as the spectrum of a reference point taken from location marked as “BG” in **Fig. 3d**. The two spectra of LD and BG are plotted together with the TAG spectrum acquired by FTIR. **Fig. 3g** shows a hyperspectral image stack (from mid-IR wavenumber of 2950 to 2700 cm^-1^ with steps of 5 cm^-1^, only 26 out of 51 images are shown for simplicity) of a zoomed-in area as highlighted in **Fig. 3d** and **Fig. 3e**, illustrating variance of the imaging contrast of lipid droplets over wavenumbers. From **Fig. 3f**,**g**, it can be observed that lipid droplets show high contrast near 2920 cm^-1^ and 2850 cm^-1^, and contrast fades out when the mid-IR wavenumber approaches 2700 cm^-1^. The two absorption peaks can be attributed, respectively, to vibration caused by symmetric and asymmetric stretching of the CH^2^ bond of triglycerides, which are the main components in lipid droplets. It is worth noting that the actual image acquisition time for this hyperspectral imaging stack (FOV: 2.5×10^5^ μm^2^, corresponding to 2.5×10^5^ pixels) with 51 frames was only 41 s (discarding data transferring time), while acquiring similar hyperspectral imaging stacks would take hours for a point-by-point scanning vibrational microscope. For instance, a hyperspectral imaging stack with FOV of 2.25×10^4^ pixels took 3.125 h when using a confocal Raman microscope.^22^

## Discussion

Here we presented phase-shifting optothermal microscopy for hyperspectral vibrational imaging of highly confluent (up to 100%) living cells in a large FOV. The unique combination of phase-shifting detection and MIR-phase generation achieves suppression of the intrinsic sample-phase and enhances sensitivity— allowing to detect small phase differences induced by vibrational absorption under wide-field mid-IR illumination. Suppression of the intrinsic sample-phase does away with the need for post-processing non-linear phase unwrapping algorithms. These algorithms are typically required in conventional quantitative-phase microscopy for the extraction of MIR-phase and their usage can result in additional imaging artifacts. In particular, with PSOM, we were able to demonstrate snapshot imaging of vibrational contrast of living cells at FOVs up to 5.18×10^5^ *µ*m^2^. Besides having a FOV that is approximately 50 times larger than current state-of-the-art (1×10^4^ *µ*m^2^; 100×100 μm^2^), PSOM achieve this large FOV without computational frame stitching^16^ and thus avoids the associated disadvantages, such as time-consumption. Imaging large numbers of cells in large FOVs as opposed to single cells in small FOVs is highly desirable in live-cell microscopy as it allows to achieve statistical relevance with a small number of quickly and easily-obtained snapshots. Additionally, benefiting from the intrinsic-phase cancellation brought about by phase-shifting— which yields sub-mrad noise-equivalent phase—we were able, for the first time, to measure highly confluent mature adipocytes (containing large organelles) as well as detect the weak vibrational signal (∼2.4 mrad) from cell membranes using PSOM. Additionally, as PSOM is able to provide hyperspectral images from large FOVs, mid-IR spectrum at each pixel in the FOV can be extracted to analyze the distribution of biomolecules of interest in a given cell population—opening up new perspectives for live-cell hyperspectral imaging and its application in label-free metabolic imaging.

As next steps, PSOM could be further developed to achieve faster live-cell hyperspectral imaging by increasing the acquisition imaging frame rate. This frame rate could be increased for instance by: 1) using a mid-IR source with a higher repetition rate than the one used here (1 kHz), so that a MIR-ON/OFF imaging cycle can be reduced from 0.5 ms down to 80 *µ*s—physical limit defined by the optothermal cycle (see **Supp. Fig. 13**); 2) implementing a polarization camera to simultaneously acquire images at 4 different analyzer angles.^23^ This will reduce the time spent on mechanical rotation of the analyzer required here and result in ability to obtain MIR-phase images in a single snapshot; and 3) application of onboard image averaging to reduce data-transfer time from camera to computer. Additionally, improvements on imaging resolution can be achieved by using a high numerical aperture (NA) objective instead of the low NA objective used in our study (NA=0.28). Furthermore, supper resolution wide-field optothermal imaging can be achieved by analyzing the time-domain optothermal signal.^24^

In summary, we foresee PSOM becoming a broadly applied imaging tool in biological research, in particular in live-cell microscopy and single-cell metabolic imaging. For example, as PSOM is a low irradiation power label-free molecular imaging modality capable of long-term longitudinal monitoring, unlike fluorescence-label based microscopy, it can continuously capture intrinsic biomolecular contrast over a long time without the risk of photobleaching. Furthermore, beyond the lipid contrast obtained here with excitation wavenumber of 2850 cm^-1^ (CH^2^), metabolism of other biomolecules, such as proteins and carbohydrates, could also be studied by extending the spectral coverage to the appropriate fingerprint region. For instance, imaging proteins using the amide bands characterized by absorption peaks between 1700 and 1400 cm^-1^ and imaging carbohydrates that can be detected in the 1300-900 cm^-1^ spectral range.

## Methods

### Experimental setup

As shown in **Fig. 1a**, the probe beam of the PSM was generated by a 532 nm pulse laser (Cobolt 06-Tor, HÜBNER GmbH & Co KG, 2 kHz), configured to emit a linear polarization beam at an orientation of 90° in relation to the x axis. A linear polarizer (LPVISC050-MP2, Thorlabs, Inc.) with a transmission axis of 45° was placed in front of the laser to obtain a 45° linearly-polarized beam. Next, the linear 45° polarized beam was divided by a birefringent beam displacer (BD27, Thorlabs, Inc.) into two orthogonally polarized beams—at 0° and 90° polarization angle with the same intensity. In the beam displacer, the 90° component (termed reference beam, **RB**) travelled straight through the beam displacer; while the 0° component (termed sample beam, **SB**) deviated 2.7 mm away from the RB path and then continued its travel parallel to RB at output of beam displacer. The two beams then passed through a half-wave plate (WPHSM05-532, Thorlabs, Inc, Fast axis: 45°), which rotated the polarization angle of both beams by 90° in relation to the fast axis of the half-wave plate. Therefore, after the half-wave plate, the polarization angle of SB and RB was 90° and 0°, respectively. The two beams then travelled through the sample plane and were, afterwards, recombined by a birefringent beam combiner (BD27, Thorlabs, Inc.). The combined beam then travelled through the objective (MY10X-803, NA=0.28; Thorlabs, Inc), which was focused on the sample plane. After the objective, the beam passed through a quarter-wave plate (WPQSM05-532, Thorlabs, Inc, Fast axis: 45°) and a rotation motor-controlled polarizer (LPVISC050-MP2, Thorlabs, Inc, termed analyzer in the main text), for phase-intensity conversion (see **Supplementary § 1**). The beam was finally focused by a tube lens to the camera (Dimax cs4, Excelitas PCO GmbH) to form a PSM image.

The pump beam was generated by a mid-infrared optical parametric oscillator (**OPO**) pulse laser (NT277, EKSPLA; 1 kHz pulse repetition rate). A germanium window was used in front of pump source to filter out the visible “signal” beam generated by the parametric process of OPO. The mid-IR (“idler” beam) was then weakly focused by a low numerical aperture gold parabolic mirror (#37248, Edmund Optics Ltd.; NA = 0.062) on the sample and the distance of parabolic mirror to the sample plane was adjusted to achieve the desirable excitation FOV.

The pump and probe laser pulses were synchronized by a trigger pulse generated by a microcontroller (Texas Instruments, MSP430F5529 LaunchPad). As shown in **Supp. Fig. 4a**, a microcontroller was connected to mid-IR laser, visible (Vis.) laser, and camera. The mid-IR laser was externally triggered by the 1kHz pulse trains to generate 1 kHz mid-IR pulses (9 ns pulse width), while the Vis. laser and camera were externally triggered by the 2 kHz pulse trains to generate 2 kHz Vis. pulses (2-3 ns pulse width), and to capture the image at the time when Vis. pulse arrived at the sample. As shown in **Supp. Fig. 4b**, since the repetition rate of the mid-IR pulse was half of the repetition rate of Vis. pulse, half of the images were acquired without mid-IR pulse excitation (MIR-OFF cycle), while the other half of images were acquired with mid-IR pulse excitation (MIR-ON cycle). The time interval between a MIR-ON image and a MIR-OFF image was 0.5 ms. Besides, a delay (termed Exposure delay or Exp. Delay) was introduced between mid-IR external trigger and Vis. external trigger so that the camera captured the image at exactly the time when the optothermal effect occurs. Varying this exposure delay results in the time-dependent optothermal transient signal as shown in **Supp. Fig. 13**, and **Supp. Fig. 14**. In the living cell measurements, we averaged 200 PSOM images to increase the CNR, which required in total 1600 PSM images for a final PSOM image (8 PSM images were required for a single PSOM image, as shown in **Fig. 1g-j**), acquired from Vis. 1600 trigger pulses within 0.8 s.

### Preparation of 3T3-L1 white adipocytes

A mouse fibroblast cell line (3T3-L1, ATCC: CRL-1658) was used as an *in vitro* model of white adipocytes. The pre-adipocytes were plated in custom-made plates (**Supp. Fig. 15b**) and cultured in growth medium -low glucose (1g l^-1^) DMEM (Merck) supplemented with 10% FBS (Merck), and 1% penicillin-streptomycin (Merck). Once 100% confluence was reached, the cells were differentiated to adipocytes. For this purpose, a differentiation medium, composed of high glucose (4.5 g l^-1^) DMEM (Merck), 10% FBS, 1% penicillin-streptomycin, 1μg ml^-1^ insulin (Sigma-Aldrich), 0.25 μM dexamethasone (Sigma-Aldrich), 0.5mM 3-isobutyl-1-methylxanthine (Sigma-Aldrich) and 1/1000 volume ABP (50 mg ml^-1^ L-ascorbate, 1mM biotin, 17 mM pantothenate (Sigma-Aldrich)) was added to the cells on day 0 and day 2. On day 4, the cells were cultured with differentiation medium supplemented only with 1μg ml^-1^ insulin and 1/1000 volume ABP. On day 6, the differentiation medium was changed back to growth medium (low glucose). Cells were kept in an incubator at 37°C with 5% CO_2_ and measured by PSOM in growth medium (low glucose). A cell-free region for the reference beam (RB) was created by covering half of dish with a medical grade tape (ARcare 90445Q, Adhesives Research, Inc.), which was removed during PSOM measurement.

### Preparation of triglyceride (TAG) drops

A 10 mg/mL suspension of TAG drops was prepared by dissolving 1 mg of 1,2-dioleoyl-3-palmitoyl-rac-glycerol (Sigma-Aldrich Inc.) in 100 μL of a chloroform–methanol solution (2:1). 10 μL of the TAG drop suspension was pipetted on a custom-made dish (**supp. Fig. 15b**, with a ZnS window as dish bottom), and left to dry at room temperature until the chloroform and methanol was completely evaporated, leaving the TAG drops on the surface of the ZnS window. After evaporation of chloroform and methanol, the dish was perfused with deionized water and covered with a cover glass, maintaining a 3 mm depth of water in the dish during the measurement. On the ZnS window, a reference region was maintained without any TAG drops for the reference beam.

### Viability test

For the viability test, we plated Hela cells in 6 custom-made dishes cultured the cells until 80% confluence was reached. The cells were divided into 2 groups: dishes 1-3 were measured by PSOM, and dishes 4-6 were used as controls. Dishes 1 -3 were measured continuously for 2 h on the PSOM stage, with one dish used per wavenumber (2850 cm^-1^, 1550 cm^-1^, or 1100 cm^-1^) tested. At these three wavenumbers, mid-IR power flux density on a sample was 0.165 *µ*W/*µ*m^2^, 0.027 *µ*W/*µ*m^2^, 0.024 *µ*W/*µ*m^2^, respectively. The corresponding control dishes (dish 4-6) were maintained in the same condition (room temperature) without mid-IR illumination. Immediately after finishing the test, to assess the negligibility of photodamage induced by PSOM, standard erythrosine B exclusion assays was performed. The cells in all the dishes (control and test) were stained with erythrosine B prior counting them and cell viability was expressed as the percentage ratio of viable irradiated cells compared to the corresponding viable non-irradiated controls. For determination of statistical significance, OriginPro9.1 software was used. Reported data corresponded to the mean ± standard deviation from three measurements.

### Contrast to noise ratio (CNR) and noise level

The contrast to noise ratio (CNR) is defined as the MIR-phase difference between the signal and the background (*φ*_*singal*_–*φ*_*BG*_) divided by the temporal noise-equivalent phase *σ*_*s*_, which takes the value of 0.26 mrad according to **Supp. Fig. 5**:

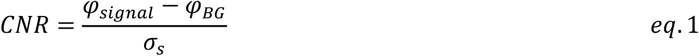

Where temporal noise-equivalent phase *σ*_*s*_ is calculated from 57 minutes of a PSOM video (**Supp. Fig. 5**) without mid-IR excitation, using:

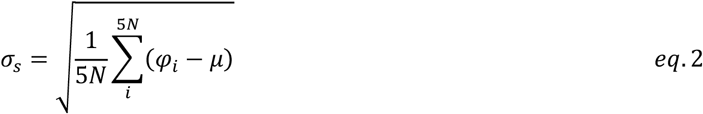

*φ*_1_ is the MIR-phase at pixel index *i* from the MIR-OFF PSOM video, and *μ* is the average of MIR-phase among 5*N* pixels, *N* is the number of frame in the video and pixels are chosen from 5 locations as shown in **Supp. Fig. 5a**. For calculating spatial noise-equivalent phase, all pixels of a single frame are used.

### Data processing

The PSOM images were processed by ImageJ. Illustration of spectra and profiles was achieved by MATLAB. For images shown in **Fig. 2g**, and **Fig. 2j**,**k**, the water absorption background (determined in cell-free region) was subtracted from the overall image, for better illustration of the cell contrast. The tracing of lipid droplets in **Supp. Fig. 11** and **Supp. Fig. 12** was achieved by TrackMate ^25, 26^.

## Acknowledgments

We thank Dr. Serene Lee for her attentive reading and improvements of the manuscript. We thank the China Scholarship Council (CSC) for financially supporting Tao Yuan’s PhD study at the Technical University of Munich.

## Author contributions

T.Y. designed and built the optical system, synchronized OPO laser and camera, automated hyperspectral imaging, and programmed the user interface for system control. L.R. provided support on polarization interferometry. F.G. provided support on biochemistry, live-cell sample preparation, and performing viability test. T.Y. performed all the rest of experiments, analyzed the results and prepared the figures. M.A.P. and T.Y wrote the manuscript. V.N. provided support on optical microscopy. M.A.P. supervised the whole study. All authors edited the manuscript.

