## Supplementary Information and Figures for "Phase-shifting mid-infrared optothermal microscopy for wide-field hyperspectral imaging of living cells"

### 1. Phase-Intensity relation

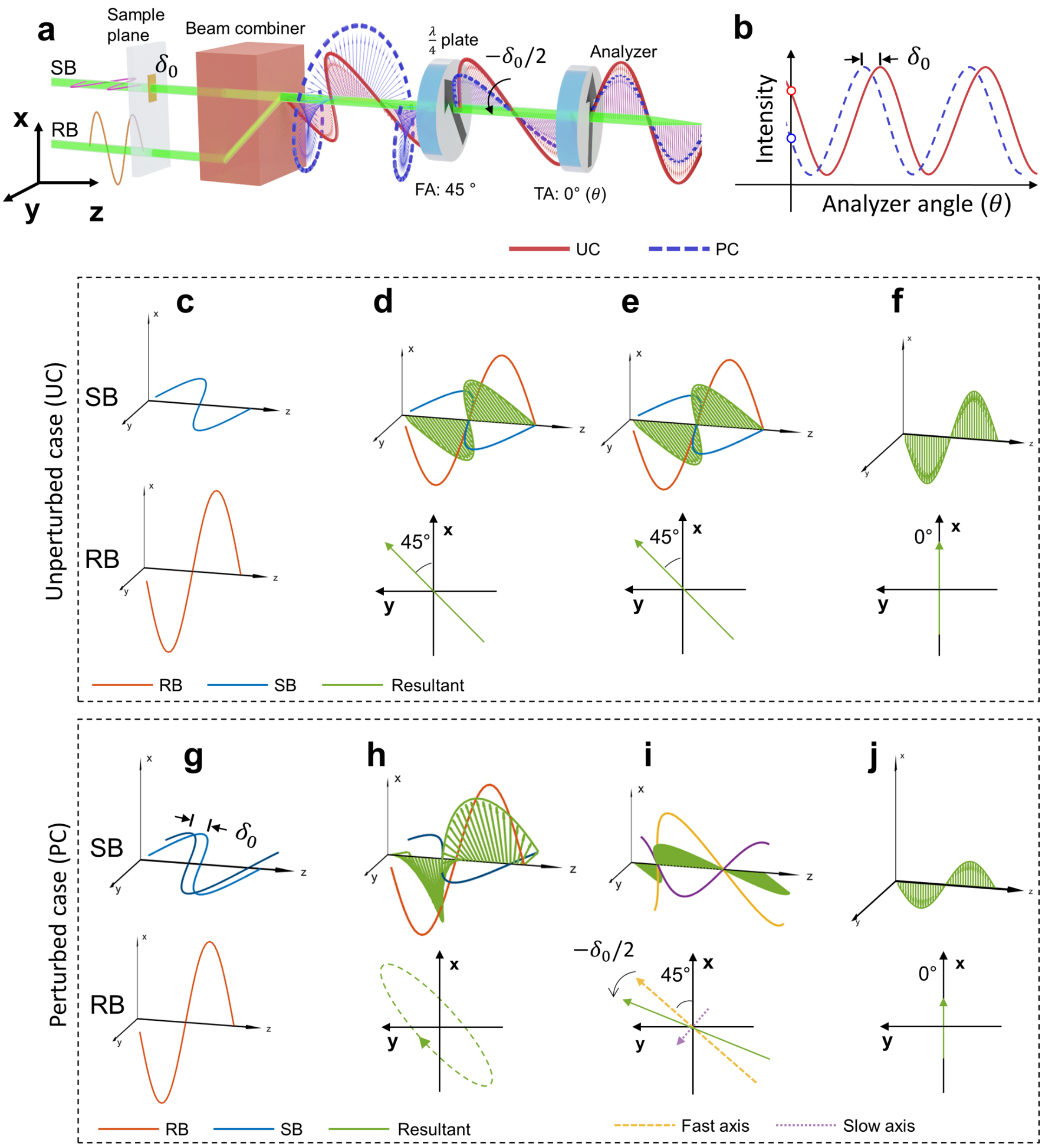


**Supp. Fig. 1. Polarization states of the probe beam as it travels through the optics. a,** 3D diagram of the system emphasizing the polarization states of probe beam in unperturbed case (UC) and perturbed case (PC). **b,** The plot of output intensity varying with analyzer angle. **c-f,** Polarization states of probe beam after (c) sample plane, (d) beam combiner, (e) quarter-wave plate, and (f) analyzer in unperturbed case. **g-j**, Polarization states of probe beam after (g) sample plane, (h) beam combiner, (i) quarter-wave plate, and (j) analyzer, in perturbed case. Second row of (d-f) and (h-j) illustrate the polarization in x-y plane. SB: Sample beam; RB: Reference beam. FA: Fast axis; TA: Transmission Axis.

**Supp. Fig. 1a,b** illustrates the polarization states of the probe-beam when interacting with the optics of the system (**Supp. Fig. 1a**), and a plot of output intensity varying with analyzer angle (**Supp. Fig. 1b**). When both beams, sample beam (SB) and reference beam (RB), keep the same optical phase (i.e., unperturbed case, see **Supp. Fig. 1c**) the combined probe beam out of the beam combiner is a 45° linearly polarized beam (**Supp. Fig. 1d**). After passing through a quarter-wave plate (fast axis: 45°, **Supp. Fig. 1a**), the combined beam keeps its 45° linear polarization state (**Supp. Fig. 1e**). If the analyzer (polarization angle can be controlled) is at 0°, only the x-axis polarization component can transmitted thought the analyzer (**Supp. Fig. 1f**), corresponding to intensity value marked by the red circle in **Supp. Fig. 1b**.

When an optical perturbation $\delta_0$ (can be the intrinsic-phase or MIR-phase) is introduced to the SB on the sample plane (i.e., perturbed case, see **Supp. Fig. 1g**), after the beam combiner, an elliptically polarized beam is the result (**Supp. Fig. 1h**). After the quarter-wave plate, this elliptical polarization beam is converted back to a linearly polarized beam (**Supp. Fig. 1i**). As shown in **Supp. Fig. 1i**, the polarization plane of the linearly polarized beam (green sinusoid wave, or green sinusoid arrow) has an angle $-\delta_0/2$ rotated relative to the unperturbed case (45°), where the minus sign stems from the fact that negative phase change (phase retardation) results in positive rotation (counterclockwise rotation). As with the unperturbed case discussed above, the analyzer (transmission angle: 0°) only allows x-axis component to pass through (**Supp. Fig. 1j**), and in this case corresponds to a lower intensity value marked by the blue circle in **Supp. Fig. 1b**. The output intensity after the analyzer follows Malus' law, which states that the intensity ($I$) of light that passes through an analyzer varies with the angle ($\theta_i$) between the light's polarization plane and the transmission axis of the analyzer, following the relationship: $I = I_0 cos^2\theta_i$ , where $I_0$ is the light intensity before entering the analyzer. Using a defined angular coordinate (defined as the clockwise angle around positive semi-axis of x), the angle $\theta_i$ can be written as $\theta_i = \pi/4 - \delta_0/2 - \theta$, where θ is the analyzer angle. When $\theta_i$ in Malus' law is substituted by this expression, we have a phase-intensity relationship:

$$I = \frac{I_0}{2}\left(1 + sin(2\theta + \delta_0)\right) \qquad eq.S1$$

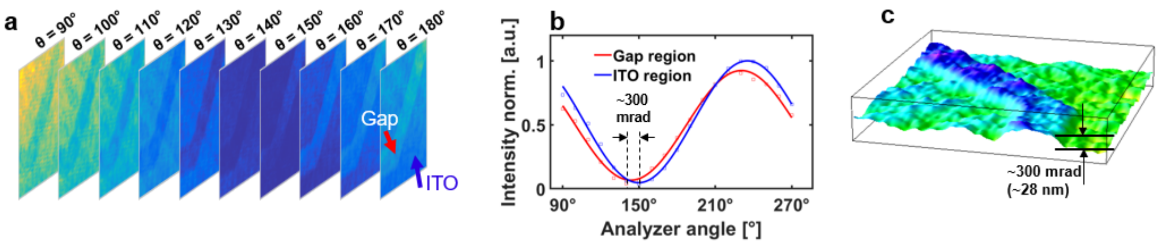


**Supp. Fig. 2. Validation of phase-intensity sinusoid relationship eq. S1 by an indium tin oxide (ITO) phase target. a**, Intensity image stack acquired by adjusting the angle of analyzer ($\theta$) from 90° to 180° with steps of 10°. Red arrow: a point on the Gap area (with absent ITO layer); Blue arrow: a point on the ITO layer.

**b,** The two points chosen from the image stack were fitted to a sinusoidal function, and the phase difference between Gap and ITO pints can be obtained as indicated. **c,** Quantitative phase image of ITO phase target constructed from the phase difference obtained at each location.

### 2. Cancellation of intrinsic-phase

For MIR-OFF images, the phase perturbation is caused by refractive index and thickness of the sample (i.e., intrinsic-phase) denoted here by $\phi_{0(x,y)}$, where x, y indicates the coordinates of the local phase perturbation in the sample plane. Following **eq. S1**, a phase-shifting module (PSM) intensity image with MIR-OFF can be written as: $I_{f(x,y)} = I_0/2\left(1 + sin\left(2\theta + \phi_{0(x,y)}\right)\right)$. When the sample is irradiated by mid-IR light (generated by pump laser as shown in **Fig. 1a** in the main text), absorption of mid-IR light by the sample creates a localized temperature change $T_{(x,y)}$, hence a local phase perturbation $\Delta\varphi_{(x,y)}$ (i.e. MIR-phase). As a result, the total phase perturbation is the sum of intrinsic-phase $\phi_{0(x,y)}$ and MIR-phase $\Delta\varphi_{(x,y)}$. Following **eq. S1**, an intensity image with MIR-ON can be written as: $I_{n(x,y)} = I_0/2\left(1 + sin\left(2\theta + \phi_{0(x,y)} + \Delta\varphi_{(x,y)}\right)\right)$. One can make a subtraction to obtain the intensity difference between MIR-ON and MIR-OFF images: $\Delta I_{(x,y)} = I_{n(x,y)} - I_{f(x,y)}$.

In PSOM, the phase-shifting mechanism requires four intensity subtraction images: $\Delta I_{1(x,y)}$, $\Delta I_{2(x,y)}$, $\Delta I_{3(x,y)}$, $\Delta I_{4(x,y)}$ from analyzer angles 315°, 0°, 45° and 90° (see **Fig. 1j** in main text). According to the phase-intensity relationship given in **eq. S1**, the four subtraction images from 4 analyzer angles (315°, 0°, 45°, or 90°) can be deduced as (after normalizing the initial intensity $I_0$ to 1, and substituting the values of θ):

$$\Delta I_{1(x,y)} = sin\left(\phi_{0(x,y)} + \frac{\Delta\varphi_{(x,y)}}{2}\right) sin\left(\frac{\Delta\varphi_{(x,y)}}{2}\right) \quad eq.\text{S2}$$

$$\Delta I_{2(x,y)} = cos\left(\phi_{0(x,y)} + \frac{\Delta\varphi_{(x,y)}}{2}\right) sin\left(\frac{\Delta\varphi_{(x,y)}}{2}\right) \quad eq.\text{S3}$$

$$\Delta I_{3(x,y)} = -sin\left(\phi_{0(x,y)} + \frac{\Delta\varphi_{(x,y)}}{2}\right) sin\left(\frac{\Delta\varphi_{(x,y)}}{2}\right) \quad eq.\text{S4}$$

$$\Delta I_{4(x,y)} = -cos\left(\phi_{0(x,y)} + \frac{\Delta\varphi_{(x,y)}}{2}\right) sin\left(\frac{\Delta\varphi_{(x,y)}}{2}\right) \quad eq.\text{S5}$$

As can be seen above, the subtraction image is expressed as the multiplication of two parts: sine or cosine function of the sum $\phi_{0(x,y)} + \frac{\Delta\varphi_{(x,y)}}{2}$, and a common term $sin\left(\frac{\Delta\varphi_{(x,y)}}{2}\right)$ shared by **eq. S2-5**. The former is dependent on intrinsic-phase $\phi_{0(x,y)}$, while the latter is independent of intrinsic-phase $\phi_{0(x,y)}$. Following the procedure described in **Fig. 1j,k**, we take the square sum of these 4 subtraction images:

$$\sigma_{(x,y)}{}^2 = \Delta I_{1(x,y)}{}^2 + \Delta I_{2(x,y)}{}^2 + \Delta I_{3(x,y)}{}^2 + \Delta I_{4(x,y)}{}^2 \qquad eq.S6$$

This results in:

$$\sigma_{(x,y)}{}^2 = 2\left[sin^2\left(\phi_{0(x,y)} + \frac{\Delta\varphi_{(x,y)}}{2}\right) + cos^2\left(\phi_{0(x,y)} + \frac{\Delta\varphi_{(x,y)}}{2}\right)\right] sin^2\left(\frac{\Delta\varphi_{(x,y)}}{2}\right) \qquad eq.S7$$

Using the formula: $sin^2(\alpha) + \cos^2(\alpha) = 1$, one can easily cancel out the dependence of intrinsic-phase $\phi_{0(x,y)}$ and obtain:

$$\sigma^2{}_{(x,y)} = 2sin^2\left(\frac{\Delta\varphi_{(x,y)}}{2}\right) \qquad eq.S8$$

Next, we apply square root to both sides:

$$\sigma_{(x,y)} = \sqrt{2}\left|sin\frac{\Delta\varphi_{(x,y)}}{2}\right| \qquad eq.S9$$

Because of the cancellation carried out in **eq. S7**, the resulting $\sigma_{(x,y)}$ in **eq. S9** is independent of the intrinsic-phase $\phi_{0(x,y)}$, which is several orders of magnitude larger than the MIR-phase. The intrinsic-phase $\phi_{0(x,y)}$ of a cell can be more than 2π rad (*1, 2*) which leads to phase wrapping artifact. In the phase wrapping artifact, all the phase ranges [2kπ, 2kπ+2π) are mapped to the range [0, 2π) (where, k = …-2,-1,0,1,2…). In the mapping relationship, the mapped phase wraps back to 0 every time the actual phase exceeds 2kπ. The phase unwrapping algorithms are usually used for recovering actual phase images, but they are error-prone and may introduce errors larger than the MIR-phase. This susceptibility to errors is even more of a challenge when dealing with images from large FOVs, where the spatial phase wrapping occurs more frequently than the pixel step of camera. The unique feature of PSOM is that it can remove the intrinsic-phase without obtaining it, circumventing the need for phase unwrapping to recover the intrinsic-phase, and making optothermal imaging of large FOVs possible. When the considered MIR-phase shift $\Delta\varphi_{(x,y)}$ is very small, we can make the following approximation from **eq. S9**:

$$\sigma_{(x,y)} \approx \left|\frac{\sqrt{2}\Delta\varphi_{(x,y)}}{2}\right| \qquad eq.S10$$

Note that the approximation can be made when the intensity images of MIR-ON/OFF are normalized to [1,0]. **Eq. S10** states that the absolute quantitative MIR-phase $\varphi_{(x,y)}$ can be obtained by multiplying $\sigma_{(x,y)}$ to a factor ($\sqrt{2}$) for construction of a PSOM image:

$$\left|\Delta\varphi_{(x,y)}\right| \approx \sqrt{2}\sigma_{(x,y)} \qquad eq.S11$$

#### 3. Relationship between the mid-IR absorption coefficient and MIR-phase

The relationship between vibrational absorption coefficient and MIR-phase $\Delta\varphi$ of a PSOM image is linearly correlated according to ref. (*3*), and the equation is as follow:

$$\frac{2\pi E}{\lambda A}\frac{(l\alpha + n\beta)}{c\rho}\mu_\omega = \Delta\varphi \qquad eq.\mathrm{S12}$$

Here, $\mu_\omega$ is the absorption coefficient at wavelength $\omega$, $E$ is the energy of a single mid-IR pulse, $\lambda$ is the wavelength of visible (Vis.) probe beam, $A$ is irradiation area of the mid-IR light and the term $\frac{2\pi E}{\lambda A}$ is system dependent. In contrast, the term $\frac{(l\alpha+n\beta)}{c\rho}$ is sample dependent, where c is specific heat capacity of sample, $\rho$ is density of the sample, $l$ is the thickness of the sample, n is the refractive index of the sample, $\alpha = \frac{dn}{dT}$ is the thermo-optic coefficient of the sample and $\beta = \frac{dl}{dT}$ is the rate of change of thickness per unit change of temperature for a sample with thickness of $l$.

For a given sample measured in a given system, $\frac{2\pi E}{\lambda A}\frac{(l\alpha+n\beta)}{c\rho}$ is a constant, and eq. S12 can be written as a linear relationship: $\kappa c\mu_\omega = \Delta\varphi$.

In summary, the MIR-phase is wavelength dependent, and the mid-IR absorption spectrum can be extracted from PSOM image stack, which acquired under different mid-IR wavelength.

### 4. Quantitative phase imaging with PSM

According to the phase-intensity formula (**eq. S1**), 4 frames of MIR-OFF intensity images at analyzer angles of 315°, 0°, 45° and 90° can be expressed as:

$$I_{1(x,y)} = I_0/2\left(1 + sin\left(630° + \phi_{0(x,y)}\right)\right) \qquad eq.\mathrm{S13}$$

$$I_{2(x,y)} = I_0/2\left(1 + sin\left(0° + \phi_{0(x,y)}\right)\right) \qquad eq.\mathrm{S14}$$

$$I_{3(x,y)} = I_0/2\left(1 + sin\left(90° + \phi_{0(x,y)}\right)\right) \qquad eq.\mathrm{S15}$$

$$I_{4(x,y)} = I_0/2\left(1 + sin\left(180° + \phi_{0(x,y)}\right)\right) \qquad eq.\mathrm{S16}$$

Tangent of intrinsic-phase can be derived using the above 4 PSM intensity images.

$$\frac{I_{2(x,y)} - I_{4(x,y)}}{I_{3(x,y)} - I_{1(x,y)}} = \frac{I_0/2\left(1 + sin\left(\phi_{0(x,y)}\right) - 1 + sin\left(\phi_{0(x,y)}\right)\right)}{I_0/2\left(1 + cos\left(\phi_{0(x,y)}\right) - 1 + cos\left(\phi_{0(x,y)}\right)\right)} = \frac{sin\left(\phi_{0(x,y)}\right)}{cos\left(\phi_{0(x,y)}\right)} = \tan\left(\phi_{0(x,y)}\right) \qquad eq.\mathrm{S17}$$

Therefore, we can obtain the quantitative phase image from these 4 PSM intensity images:

$$\phi_{0(x,y)} = tan^{-1}\left(\frac{I_{2(x,y)} - I_{4(x,y)}}{I_{3(x,y)} - I_{1(x,y)}}\right) \qquad eq.S18$$

### 5. Figures

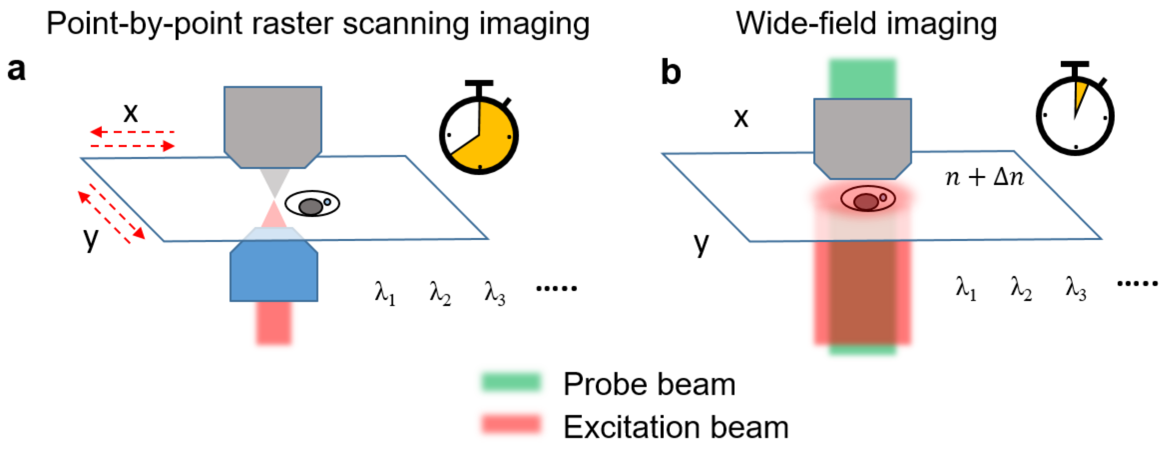


**Supp. Fig. 3. Comparison of point-by-point raster scanning imaging and wide-field imaging. a,** In point-by-point scanning imaging, the excitation beam is focused into a diffraction limit spot, and an image is constructed from data acquired at each point in the FOV. Due to the scanning mechanism, hyperspectral imaging from multiple excitation wavelengths ($\lambda_1, \lambda_2, \lambda_3 \ldots$) takes long time. **b,** In wide-field imaging, a broad mid-IR beam is used for excitation. Visualization of a sample in a large FOV can be achieved quickly via capturing a snapshot of detected optothermal-related phase change.

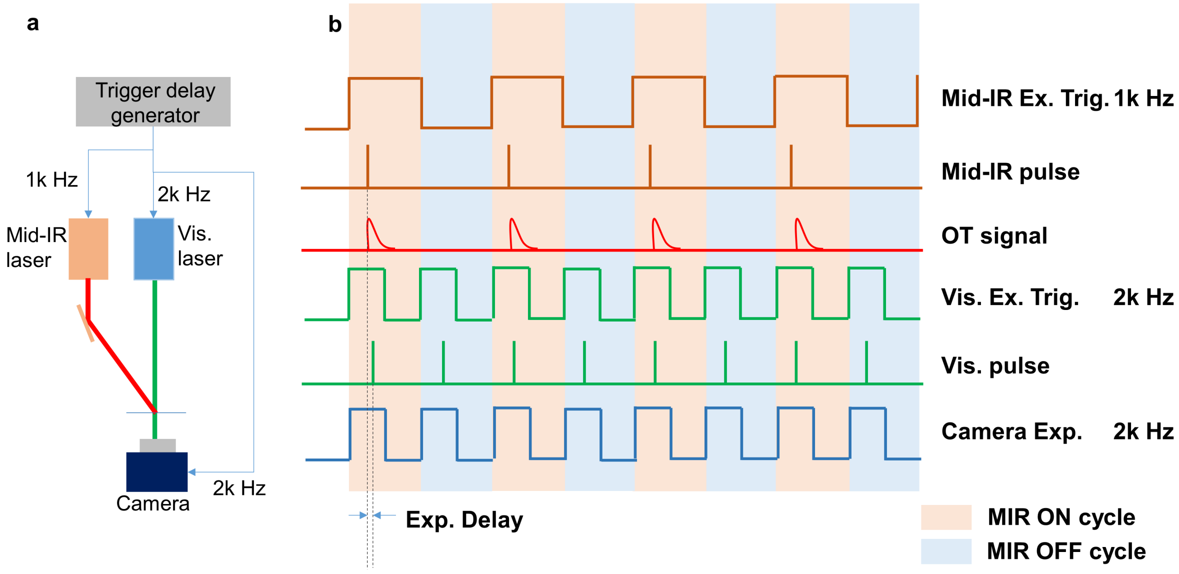


**Supp. Fig. 4. System synchronization**. **a**, Diagram of a simplified PSOM, emphasizing the synchronization connection and pump-probe mechanism. **b,** Diagram of the trigger pulse trains and their corresponding mid-IR/Vis. pulse outputs, and optothermal (OT) signals. Mid-IR Ex. Trig.: External trigger for the mid-infrared optical parametric oscillator (OPO). Vis. Ex. Trig.: External trigger for the 532 nm pulse laser. Camera Exp.: External trigger for camera exposure. The three synchronized trigger signals (mid-IR Ex. Trig., Vis. Ex. Trig.

and Camera Exp.) are generated by a trigger delay generator, where the mid-IR Ex. Trig. is a 1k Hz square pulse train, both Vis. Ex. Trig. and Camera Exp. are 2k Hz square pulse trains. As demonstrated in (b), the camera operates with a frame rate of 2k Hz, triggered by the 2k Hz signal; the exposure cycle with mid-IR pulse is called the MIR ON cycle, while the cycle without mid-IR pulse is called the MIR OFF cycle, the corresponding images are referred to as MIR-ON images and MIR-OFF images, respectively. An exposure delay (Exp. Delay) is strategically introduced to the two 2k Hz signals (Vis. Ex. Trig. and Camera Exp.), so that the mid-IR pulse and Vis. pulse reach the sample at the same time. Time-dependent optothermal transient signal can be obtained by varying the exposure delay.

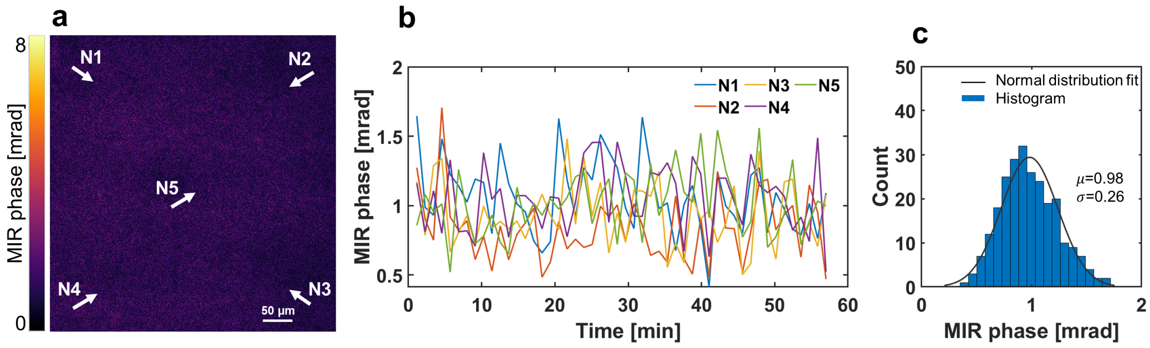


**Supp. Fig. 5. Temporal noise measurement. a**, A PSOM image acquired without mid-IR irradiating the FOV. **b,** Temporally change of MIR-phase from 5 locations of the FOV, as marked in (a). Measurement duration: 57 minutes. **c,** Histogram of all the temporal data points from (b), and the corresponding normal distribution fit curve. $\mu$: mean; $\sigma$: standard deviation. Temporal noise-equivalent phase of 0.26 mrad is obtained (See **Methods** of main text), corresponding to equivalent optical path length of 22 pm (cosidering visible laser wavelenght of 532 nm).

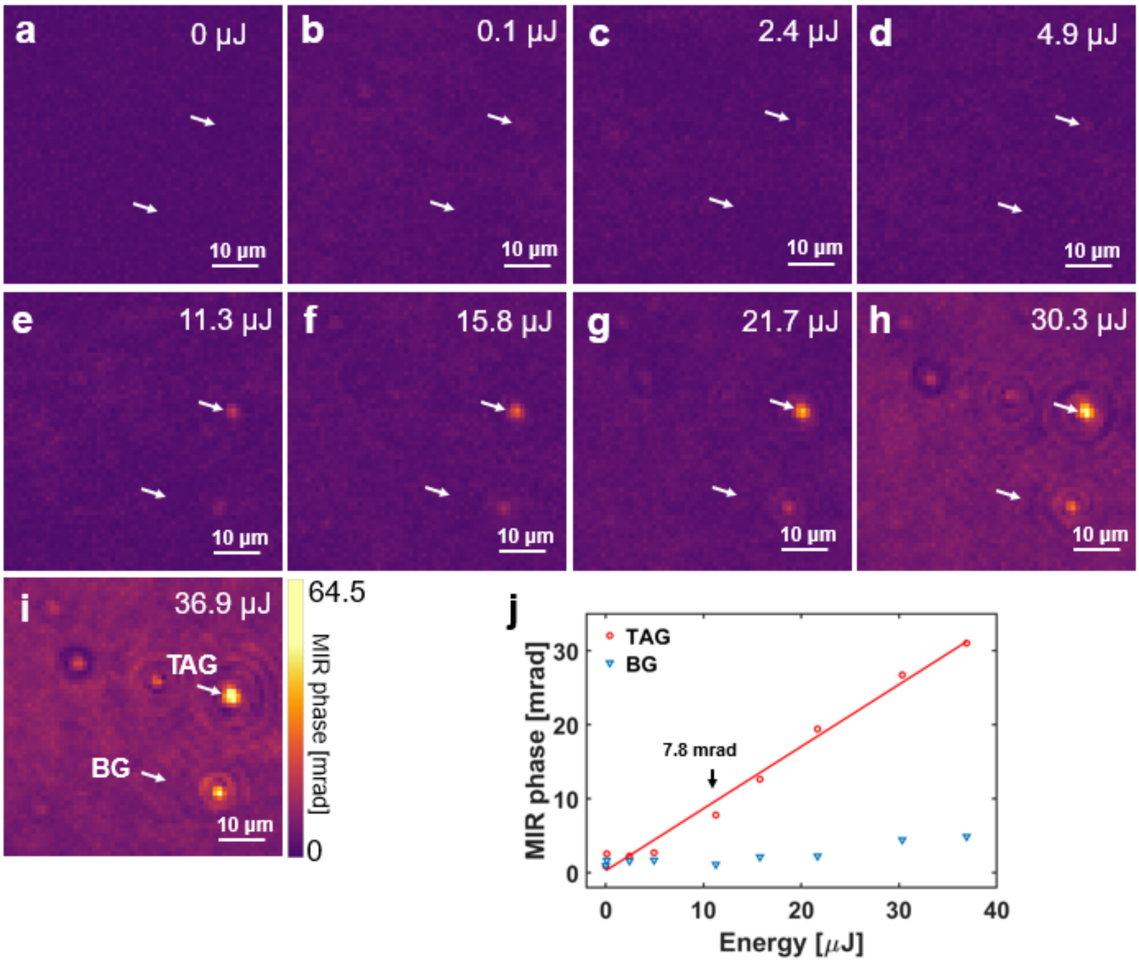


**Supp. Fig. 6. Linear relationship between mid-IR pulse energy and MIR-phase of PSOM image. a-i**, PSOM images with different mid-IR pulse energy directed on the sample, ranging from 0 μJ to 36.9 μJ. The visibility of synthetic triglyceride (TAG) drops increases as the pulse energy increases from (a) to (i). **j,** Plot of the MIR-phase values (averaged over a 3×3 pixel area) at a synthetic TAG drop and in an area selected for acquiring background (BG) values. The MIR-phase of a particular TAG drop (indicated in Fig. 5i) is linearly correlated with excitation energy. In (e), under mid-IR pulse energy of 11.3 μJ/pulse (corresponding to power flux density of 0.05 $\mu W/\mu m^2$), the TAG drop can be distinguished with a contrast-to-noise ratio (CNR) of 26:1 (MIR-phase of 7.8 mrad).

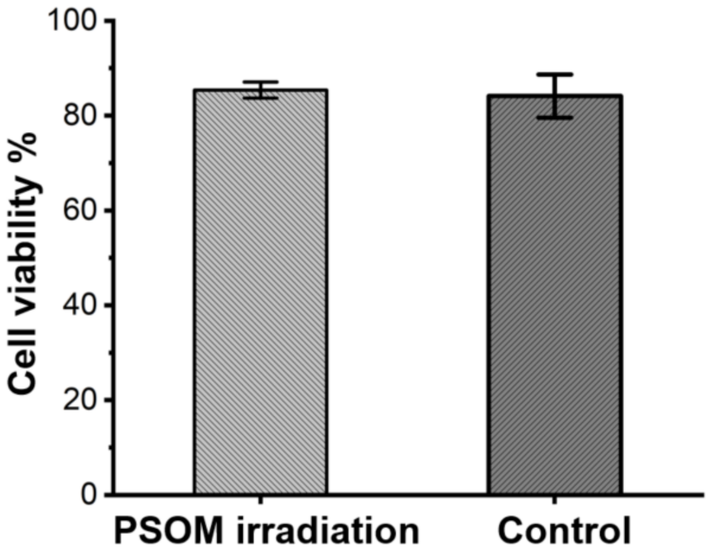


**Supp. Fig. 7. Viability assessment of HeLa cells with and without PSOM irradiation.** In this test Hela cells were imaged continuously for 2 h using PSOM. The test was performed 3 times on 3 different dishes for each condition (with or without irradiation). The controls were prepared and maintained in the same condition (room temperature) as the imaged sample and were not subjected to mid-IR excitation. The bars represent the mean cell viability of PSOM irradiated (85.4%) samples ($N = 3$) and control (83.8%). The error bars represent the standard deviation (PSOM irradiated samples: 3.9%; control: 8.2%).

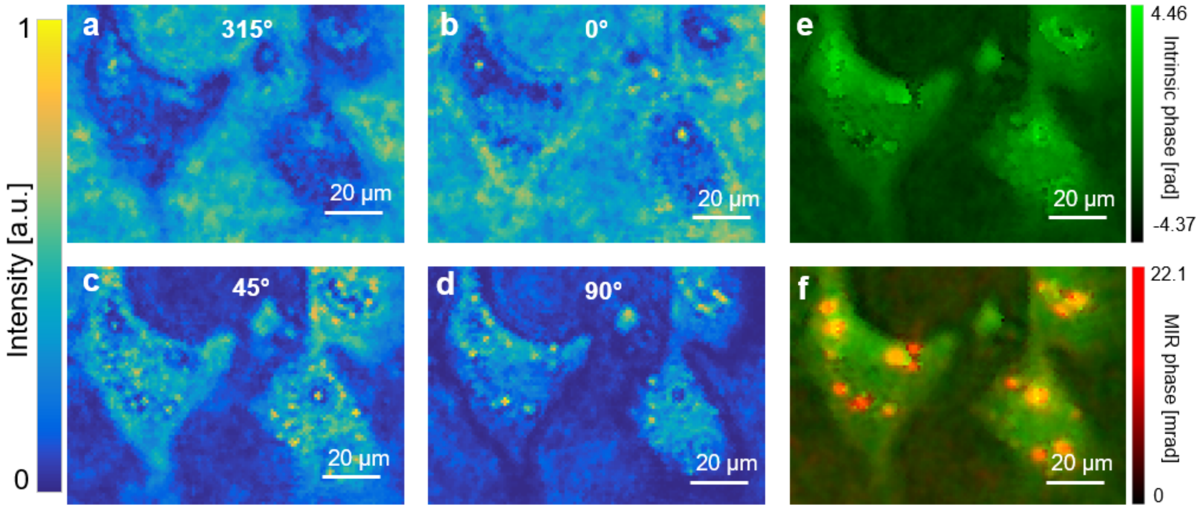


**Supp. Fig. 8. Intensity images and quantitative intrinsic-phase of 3T3-L1 cells (same FOV as Fig. 2g,h). a-d**, MIR-OFF intensity images of cells at analyzer angles of 315°, 0°, 45°, and 90°. **e**, Quantitative intrinsic-phase reconstructed from (a-d) using eq. S18. The quality guide phase unwrapping method was used for reconstructing the actual intrinsic-phase (*4, 5*). **f**, Merge of quantitative intrinsic-phase image and MIR-phase image at 2850 $cm^{-1}$.

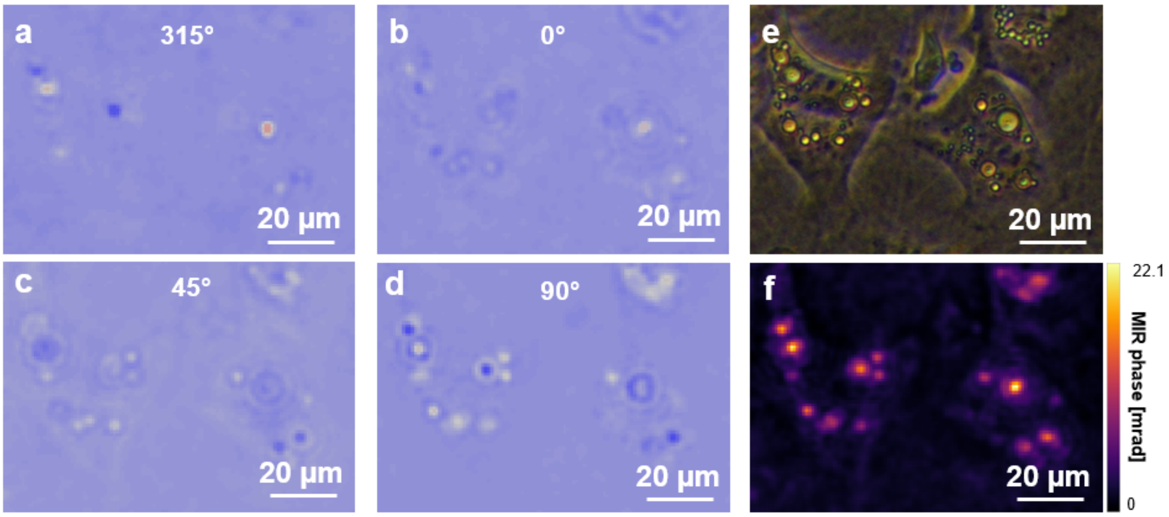


**Supp. Fig. 9. Intensity subtraction images from measurements on live cells at mid-IR wavenumber of 2850 cm$^{-1}$. a-d**, Subtraction images for analyzer angles of 315°, 0°, 45°, and 90°. **e**, Image acquired by Zernike's Phase Contrast (ZPC) microscope of the same FOV as (a-d). **f,** PSOM image constructed using eq. S11.

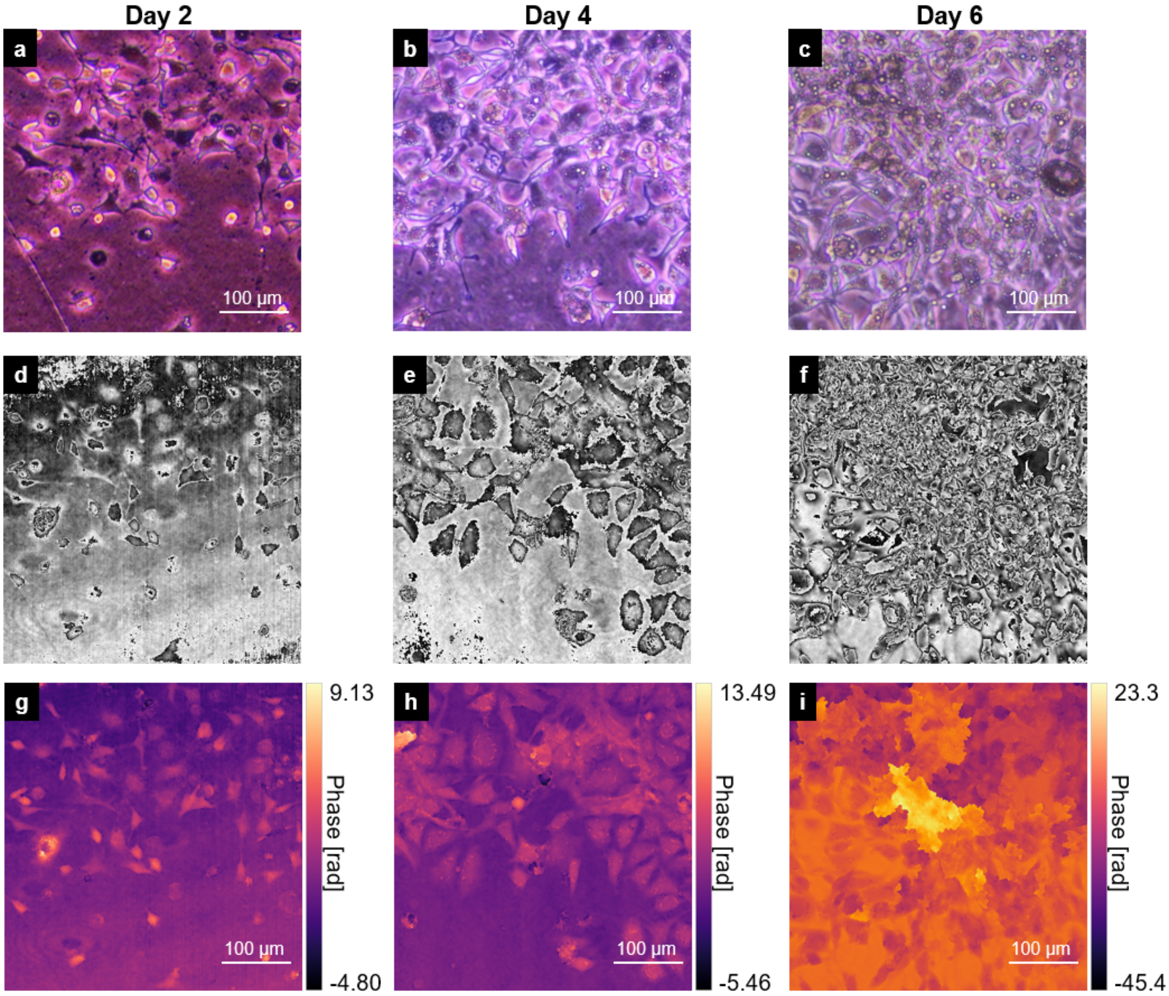


**Supp. Fig. 10. Intrinsic-phase of 3T3-L1 cells on days 2, 4, and 6 after differentiation. a-c,** ZPC micrographs. **d-f,** Wrapped phase for the FOVs corresponding to (a-c) (applying eq. S18). **g-i,** Full range intrinsic-phase images reconstructed using quality guide phase unwrapping method (*4, 5*). At high confluence (i), due to strong scattering caused by clumps of cells, the phase unwrapping method failed to recover morphological information.

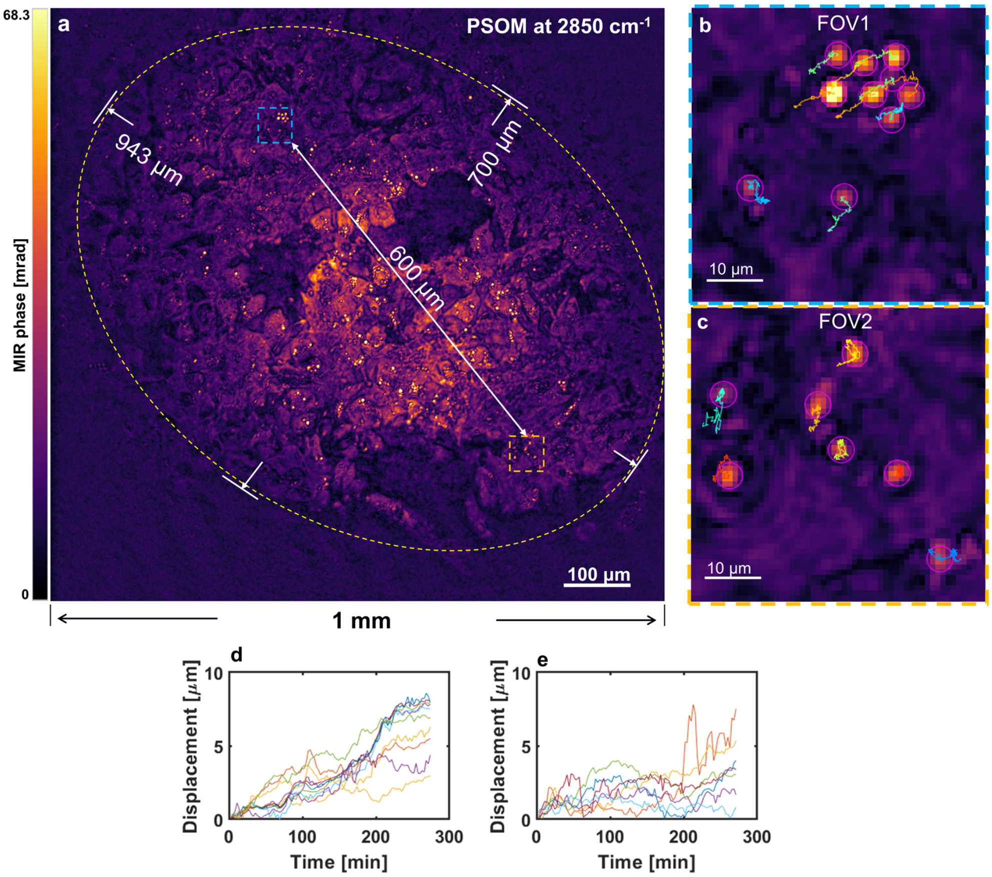


**Supp. Fig. 11. Live-cell imaging on large FOV. a,** Large FOV attained by expanding the mid-IR illumination spot used to measure dynamic lipid droplets (LDs) in living cells (3T3-L1). The effective FOV is highlighted by the dashed ellipse with major axis of 943 µm, and minor axis of 700 µm, corresponding to an FOV of $5.18\times10^5$ µm$^2$. The time taken to capture this image is 0.8 s. **b,c,** Two zoomed in regions in FOV from (a), illustrating the paths of LDs tracked during a monitoring duration of 4.5 h. **d,e,** Plots of the LDs' displacement from their original locations, obtained from tracked LDs in (b) and (c) respectively.

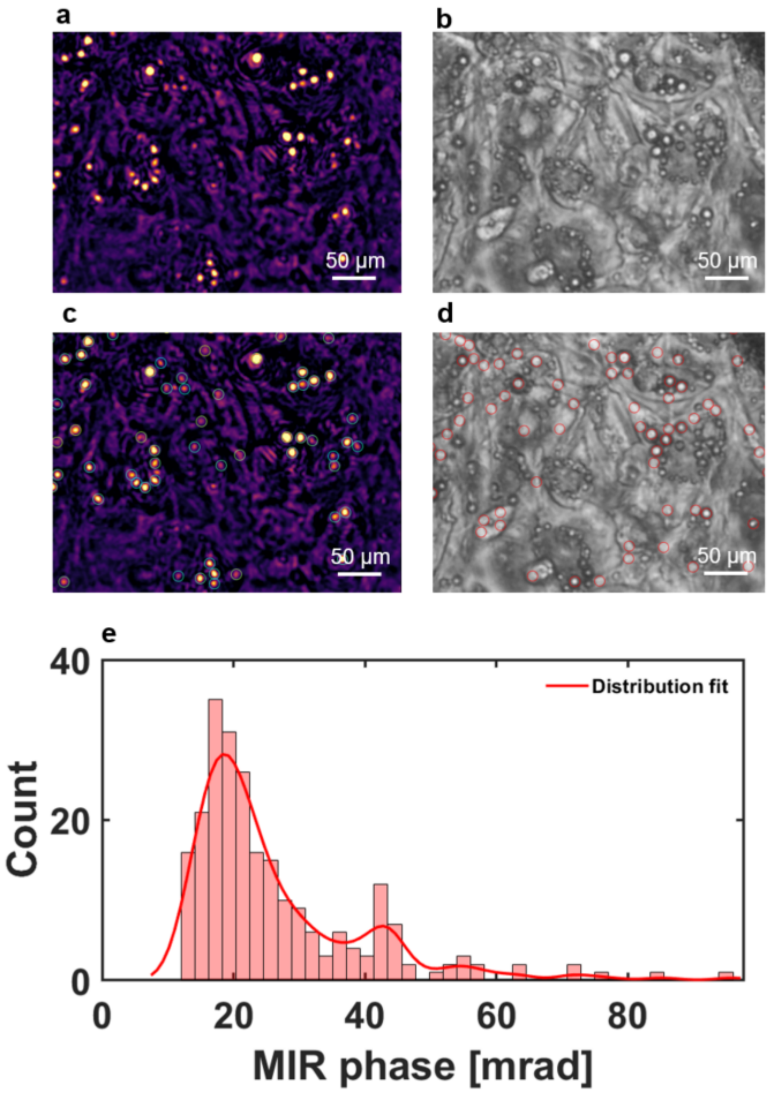


**Supp. Fig. 12. Lipid droplet segmentations on PSOM images and ZPC images. a,b,** Zoomed in images of the dashed rectangles in the FOV of Fig. 3a,b. **c,d,** The detection program, TrackMate (6, 7), automatically marks lipid droplets with circles. Using the same method, LDs in the PSOM micrograph (c) can be easily segmented from background, while LDs in the ZPC micrograph (d) are not segmented correctly due to strong phase background. **e,** Statistic analysis of the segmented lipid droplets obtained from the whole FOV of Fig. 3b, illustrating their MIR-phase distribution.

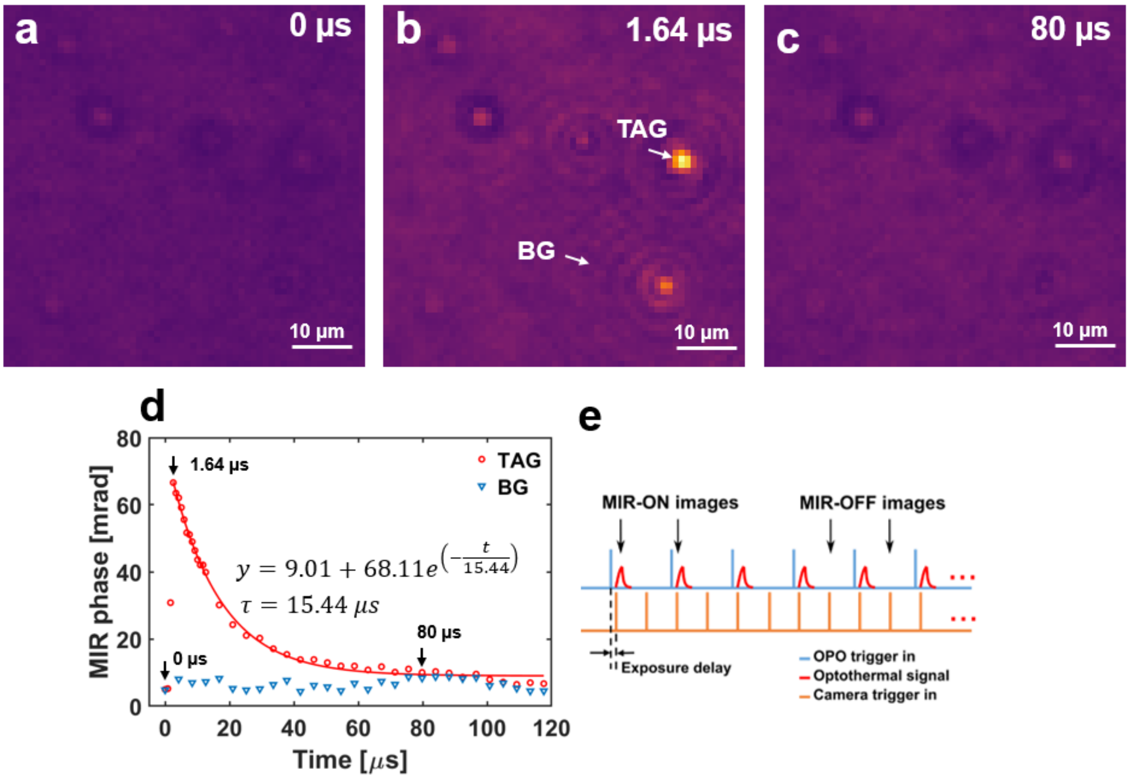


**Supp. Fig. 13. Time-dependent optothermal transient signal measured from synthetic TAG phantom. a-c,** PSOM image of the FOV at exposure delay of 0 µs, 1.64 µs and 80 µs. **d,** Plot of MIR-phase values measured at a TAG drop and an area defined as background (BG) as marked in (b). **e,** The trigger pulse train used in this measurement. OPO: optical parametric oscillator.

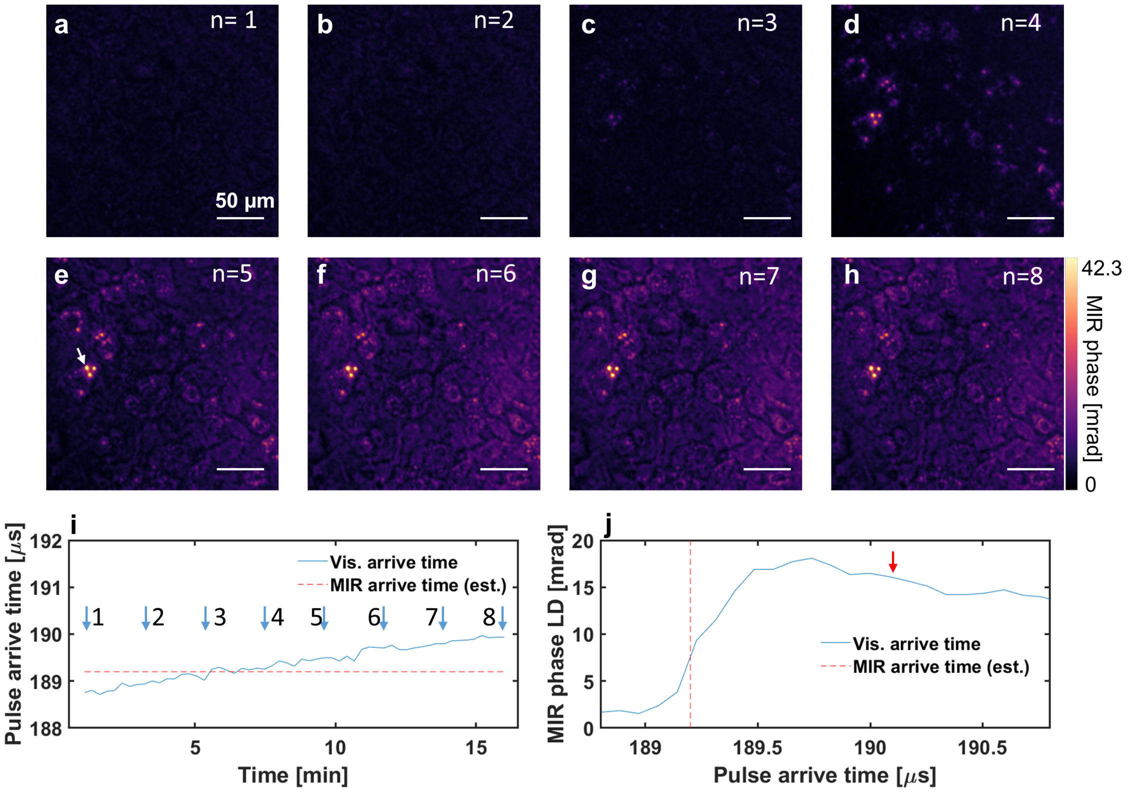


**Supp. Fig. 14. Time-dependent optothermal transient signals measured from live cells**. **a-h**, PSOM images acquired by varying the exposure delays (see **Method**, and **Supp. Fig. 4**). **i**, The Vis. pulse arrival time measured by a photodiode detector and a synchronized data acquisition card (DAQ). The Vis. pulse arrival time for corresponding figures of (a-h) is marked by a number "1" to "8". **j,** Plot of MIR-phase vs. time for the LD marked in (e). Vis. arrive time, and MIR arrive time are defined as the time taken for the Vis. pulse to hit the sample plane, and the time taken for the mid-IR pulse to hit the sample plane after triggering the mid-IR . Vis.: visible laser; MIR: mid-infrared laser.

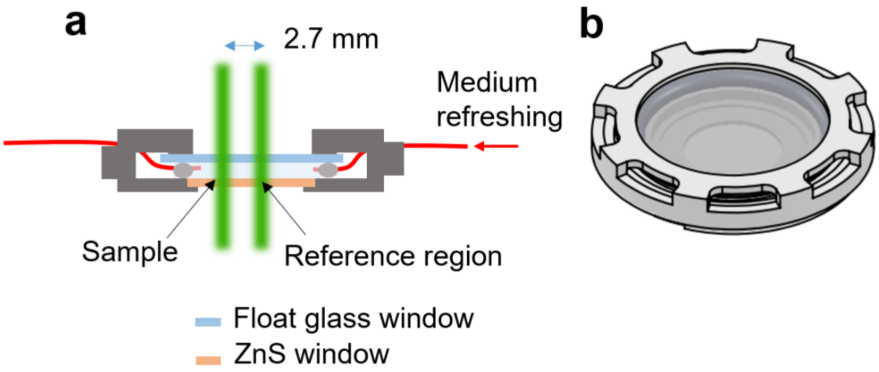


**Supp. Fig. 15. Configuration for measuring living cells**. **a**, Side view of the dish during the measurement of living cells. **b,** 3D view of the dish. The cells are maintained in a dish with two parallel windows (as see in Fig. a, bottom: ZnS window; top: Float glass window). The two probe beams 2.7 mm apart pass through the dish, with one beam passing through a sample region (with cells) and the other beam passing through a reference region (with no cells). For monitoring over a long duration, the medium can be refreshed through an inlet going into the dish.

**Supp. Table 1.** Comparison of PSOM with state-of-the-art optothermal/photothermal microscopes.

| | **WOMiM** (*8*) | **BSTP** (*3*) | **ADRIFT** (*9*) | **BS-IDT** (*10*) | **PSOM** |
|---|---|---|---|---|---|
| FOV (μm$^2$) | 2.5 ×$10^4$ | ~2.0×$10^3$ | ~2.0×$10^3$ | 1.0×$10^4$ | 5.2 ×$10^5$ |
| MIR shot-noise (mrad) | Not provided (-) | ~5 | ~ 1 | Not Applicable (NA) | 0.33 |
| SNR | ~50 | 30-100 | - | - | 203 |
| Excitation irradiance/PFD (μW/μm$^2$) | 0.39 | 30 | - | ~0.5 | 0.05 |
| Resolution (μm) | 1.85 | 0.96 | - | 0.35 | 2.03 |
| Imaging speed (Frame per second) | 50 | 1-100 | 5 | 0.05-6 | 0.017 |
| High Confluence (Yes/No, y/n)? | n | n | n | n | y |
| Phase-wrapping artifacts (y/n)? | y | y | y | NA | n |
| Demonstrated cell viability (y/n)? | n | n | n | n | y |
| Living cells (y/n)? | n | y | y | n | y |

**Supp. Table 2.** Comparison of focal excitation and wide-field excitation imaging methods. (*3, 11-13*)

| | **Focal excitation** | | **Wide-field excitation** |
|---|---|---|---|
| | Galvo scanning | Mechanical-scanning | |
| Has trade-off between imaging Time & FOV? | Yes | Yes | No |
| Excitation irradiance μW/μm$^2$) | 1.5×$10^4$ | 8.0 | 0.05 |
| Imaging speed (fps) | ~10-30 (For FOV of c.a. 1×$10^6$ μm$^2$) | 0.0055 (For FOV of c.a. 1×$10^6$ μm$^2$) | 100 (Does not depend on FOV) |
